# The repertoire of short tandem repeats across the tree of life

**DOI:** 10.1101/2024.08.08.607201

**Authors:** Nikol Chantzi, Ilias Georgakopoulos-Soares

## Abstract

Short tandem repeats (STRs) are widespread, dynamic repetitive elements with a number of biological functions and relevance to human diseases. However, their prevalence across taxa remains poorly characterized. Here we examined the impact of STRs in the genomes of 117,253 organisms spanning the tree of life. We find that there are large differences in the frequencies of STRs between organismal genomes and these differences are largely driven by the taxonomic group an organism belongs to. Using simulated genomes, we find that on average there is no enrichment of STRs in bacterial and archaeal genomes, suggesting that these genomes are not particularly repetitive. In contrast, we find that eukaryotic genomes are orders of magnitude more repetitive than expected. STRs are preferentially located at functional loci at specific taxa. Finally, we utilize the recently completed Telomere-to-Telomere genomes of human and other great apes, and find that STRs are highly abundant and variable between primate species, particularly in peri/centromeric regions. We conclude that STRs have expanded in eukaryotic and viral lineages and not in archaea or bacteria, resulting in large discrepancies in genomic composition.

## Introduction

The rapid decline in the cost of DNA sequencing has facilitated the generation of complete genomes for an ever growing number of organisms, across the tree of life. A number of large, international consortia are currently underway with the goal of sequencing genomes representing the diversity found in nature (Lewin et al. 2018; Darwin Tree of Life Project Consortium 2022; Land et al. 2015). This offers a remarkable opportunity to increase our understanding of genome architecture, evolution, and diversity in unprecedented depth and breadth. Additionally, the recent completion of the human genome through the Telomere-to-Telomere (T2T) Consortium (Nurk et al. 2022) enables the examination of the distribution and frequency of repeat elements in regions of the genome that were only partially annotated previously, including centromeres and telomeres. Another recent effort includes the T2T-Primates consortium (Hoyt et al. 2022), which has completed the generation of diploid assemblies for multiple non-human primate species and through which we can gain insights into the diversity, evolution and plasticity of different repeats in the primate lineage.

Repetitive DNA sequences are widespread in the genomes of various life forms. For example, over half of the human genome is composed of repetitive sequences (Lander et al. 2001). Nevertheless, we still do not fully understand the landscape of repeat elements in organisms across the tree of life. Initially thought of as junk DNA, repeat elements are fast evolving, and are usually classified into two major classes, tandem repeats and dispersed repeats, the second of which also encompasses transposable elements (Paço, Freitas, and Vieira-da-Silva 2019). Short tandem repeats (STRs) are defined as multiple consecutive copies of an oligonucleotide repeat unit, often defined for up to 1-9 base-pairs (bps) unit length (Cer et al. 2013). These are highly polymorphic, representing a major source of genetic variation, due to their heightened mutation rate (Sun et al. 2012).

In the human genome there are more than one million tandem repeats (Hannan 2018), which vary substantially in the population (Shi et al. 2023; Liao et al. 2023; Tanudisastro et al. 2024). STRs are mutational hotspots, associated with genomic instability (Khristich and Mirkin 2020), providing a source for genomic diversity, polymorphism and organismal adaptation, while they are also linked to a number of human diseases (Mirkin 2007; Hannan 2018). STRs trigger slippage events during DNA replication, leading to frequent mutations in the number of repeat units, through insertions and deletions (Murat, Guilbaud, and Sale 2020). As a result of the genomic instability associated with them, STRs are causative for a number of human diseases (Mirkin 2007), including Mendelian disorders (La Spada and Taylor 2010), neurodegenerative disorders (Hannan 2018), and different cancer types (Erwin et al. 2023), and contribute to multiple complex traits and phenotypes (Hannan 2018; Gall-Duncan et al. 2022). In microorganisms, STRs are also known to be related to pathogenicity and genomic variability (Jackson et al. 1997; Field and Wills 1998).

Furthermore, STRs can form non-canonical DNA structures such as Z-DNA, G-quadruplexes and hairpins, which are also associated with higher mutation rate and have a plethora of functional roles (Georgakopoulos-Soares et al. 2018; Georgakopoulos-Soares, Parada, and Hemberg 2022; Georgakopoulos-Soares et al. 2022; Guiblet et al. 2021; Wang and Vasquez 2023). STRs have been linked to a number of functions, including roles in gene regulation (Bakhtiari et al. 2021; Fotsing et al. 2019; Liang et al. 2015; Horton et al. 2023), recombination (Treco and Arnheim 1986; Wahls, Wallace, and Moore 1990), epigenetic changes (Quilez et al. 2016) and genome architecture (Farré et al. 2011) among others. The instability of STRs have various implications and applications in molecular biology. They enable DNA forensics, PCR-based genotyping and phenotyping, fine-scale phylogenetic analysis, paternity testing, linkage-disequilibrium mapping, and hitchhiking mapping (Fungtammasan et al. 2015; Wyner, Barash, and McNevin 2020). In eukaryotic organisms, STRs are also highly prevalent in pericentromeric and centromeric regions (Patchigolla and Mellone 2022; Melters et al. 2013). However, systematic study of these regions has been challenging due to the limitations of short-read sequencing technologies, which struggle to accurately determine the sequence composition of these parts of the genome (Need and Goldstein 2009).

Because of their higher mutation and turnover rates, STRs are of extreme interest to understand evolution. STRs play a crucial role in driving species evolution and contribute significantly to the diversity and adaptability of life forms (Zhou, Aertsen, and Michiels 2014). Multiple studies have investigated the prevalence of STRs in specific taxonomic groups (Srivastava et al. 2019; Ding et al. 2017; Song et al. 2021; Yuan et al. 2018; Lin and Kussell 2012; Mrázek, Guo, and Shah 2007). Nevertheless, despite their importance, there is currently no comprehensive analysis of the distribution, diversity and topography of STRs among taxa spanning the different domains, kingdoms and phyla in the tree of life.

Here we perform the largest comprehensive examination of tandem repeat variations across 117,253 organismal genomes spanning all major taxonomies of the tree of life. We find that STRs are most enriched in eukaryotic organisms, with *Plasmodium falciparum* exhibiting the highest STR density. Using genome simulation experiments we find that on average archaea and bacteria show a relative depletion of STRs, whereas eukaryotes show the highest STR enrichment (roughly 10-fold), relative to what is expected by chance. We find large differences in the frequency of STRs with different repeat unit lengths, which are also influenced by the taxonomic group and the genomic compartment. Finally, we perform an in depth analysis of the recently published T2T primate genomes. We find marked enrichment of STRs in pericentromeric and centromeric regions, which have rapidly and dynamically evolved in primate genomes. These enrichments are driven by specific STR units and tend to be dinucleotide and pentanucleotide. Surprisingly, large differences in the STR composition, frequency and location are also observed between chromosomes in these genomes. These findings indicate large differences in the frequency and topography across the tree of life, acting as a highly dynamic, and polymorphic genomic element.

## Results

### Systematic identification of short tandem repeats in organismal genomes

We examined 117,253 organismal genomes spanning the tree of life for the genome-wide prevalence of short tandem repeats (STRs). STRs were defined for unit lengths between one and nine base-pairs (bps). We identified STR sequences across the genomes of each available organism, creating a comprehensive genome-wide dataset with species from diverse taxa across the tree of life. In total, we identified 81,864,637 STRs across the 117,253 organismal genomes. We observed that the average STR density was 1.51 STR bps per kB (**Supplementary Figure 1**). However, we find that there are large differences depending on the organism and its taxonomic group. Across all the examined species we observe that *Plasmodium falciparum* shows the highest STR genomic density (109.18 bps/kB), with a 72.3-fold enrichment over the average STR density across organismal genomes, whereas 19,712 viruses and 5 bacteria lack STRs altogether (**Supplementary Figure 1**). This is consistent with a previous work studying 719 organismal genomes, in which *Plasmodium falciparum* was also found to have the highest STR density (Srivastava et al. 2019) and consistent with it having a highly repetitive, AT-rich genome (Zilversmit et al. 2010).

Next we investigated if there is a correlation between the genome size and the density of STRs in each of the three domains of life and viruses. We find that archaea and eukaryotes display positive correlations (Spearman correlations of R=0.44 and R=0.39), whereas bacteria show a negative correlation (Spearman correlations of R=-0.26) (**Figure 1a**). When comparing the domains of life, the highest genomic density of STRs was observed in eukaryotes with 581.42 STR bps per megabase (Mb), whereas the lowest was observed in archaea with 75.07 STR bps per Mb (**Figure 1b**; Mann-Whitney U test, p-value<0.0001). For viruses, we further subdivided the analysis by their host type and found that viruses of eukaryotic hosts had the highest genomic density of STRs (**Figure 1b**; Mann-Whitney U test, p-value<0.0001). These findings indicate that STRs are most abundant in eukaryotic genomes, in which their frequency correlates with genome size.

**Figure 1:**
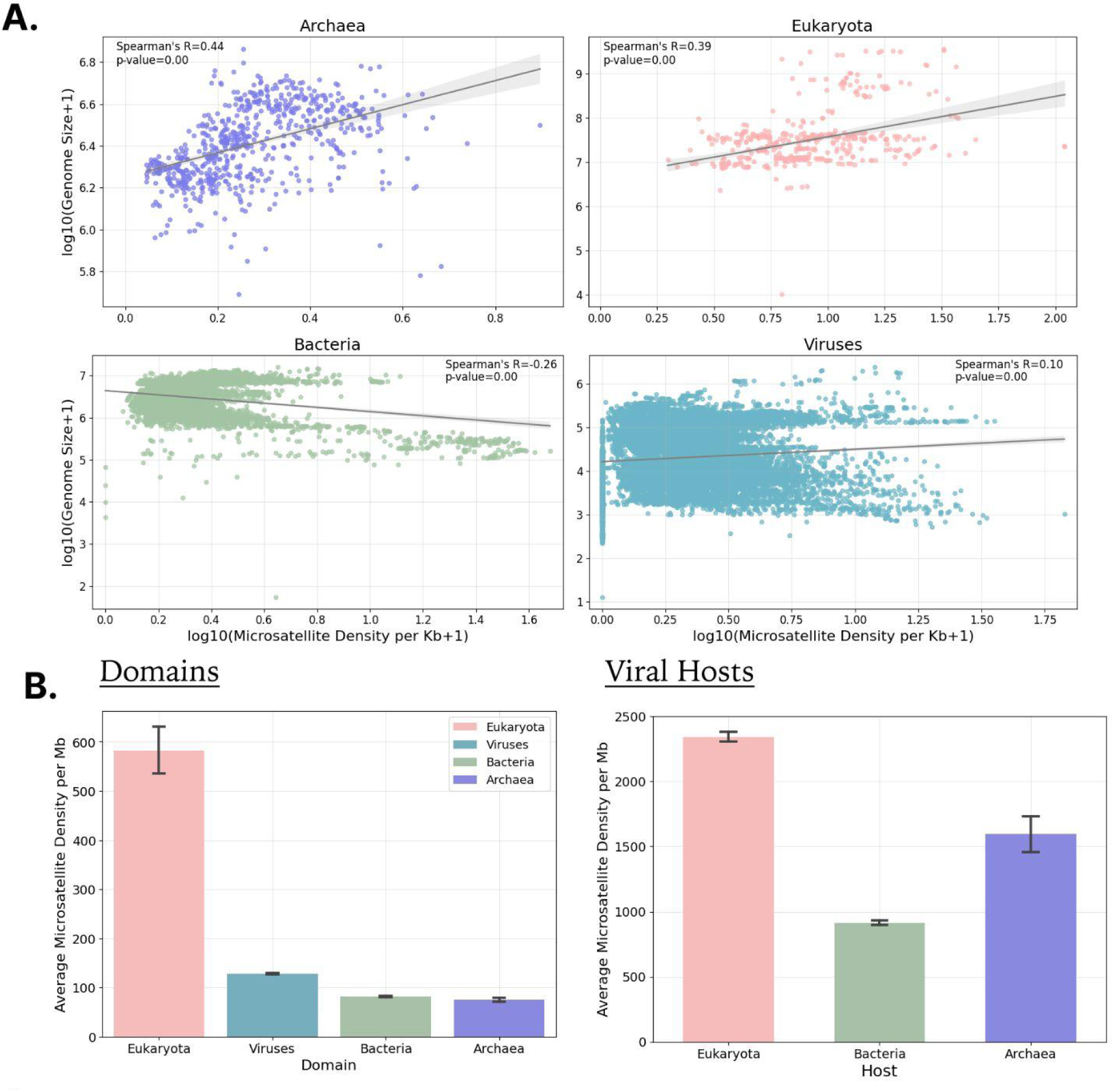
Characterization of STRs in 117,253 organismal genomes across the three domains of life and viruses. **A.** Association between the genome size and the proportion of the genome covered by STRs, presented separately for the three domains of life and viruses. d. Representation of the number of shared sequence arms for STRs identified in organismal genomes across the three domains of life and viruses. **B.** Average STR bp density per kB for organisms in the three domains of life and viruses and for viruses relative to the viral host domain. Error bars represent standard error.

### Characterization of tandem repeat frequencies across taxa

Next, we examined differences in the distribution and density of STRs between taxonomic subgroups. We observe that at the kingdom level, Protista, Animalia, Plantae and Fungi, all of which are eukaryotic, show the highest STR density, with Protista having an average of 1,425.12 STR bps per Mb. This is followed by different viral kingdoms, whereas Eubacteria and Archaebacteria show the lowest genomic STR density (**Figure 2a**), consistent with our observations at the domain level. We also examined the average STR density across organisms belonging to the same phylum. We find that consistently eukaryotic phyla show the highest STR density, with Euglenozoa and Apicomplexa having 1,550.79 and 1,442.33 STR bps per Mb (**Figure 2b**).

**Figure 2:**
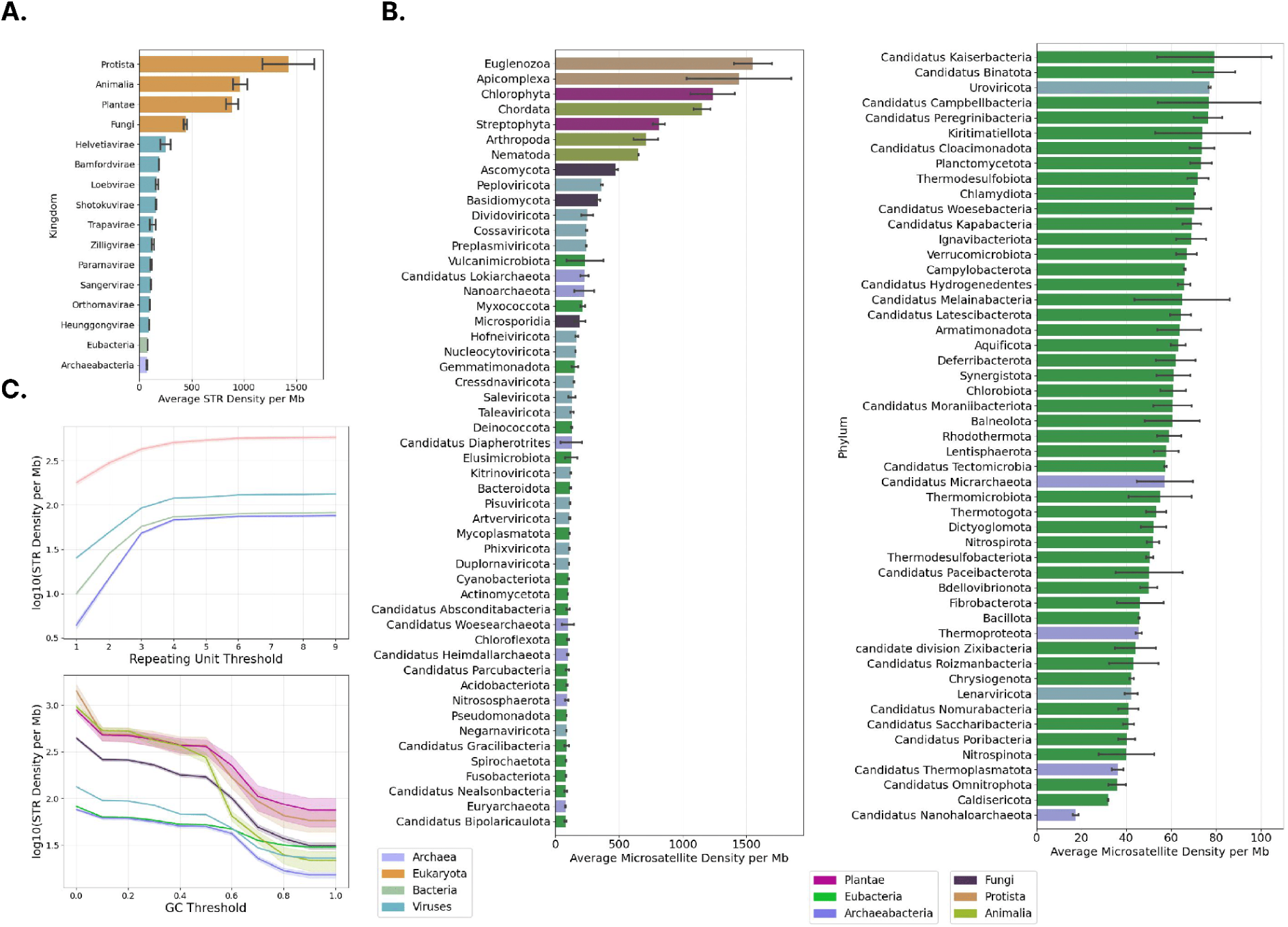
Taxonomic characterization of STRs across the tree of life. **A.** Density of STRs in each kingdom. Error bars show standard error. Coloring is performed at the domain of life level. **B.** Density of STRs in each phylum. Error bars show standard error. Coloring is performed at the kingdom level. **C.** STR density relative to the GC content threshold in the STR sequence. Results shown at the kingdom level and the domain level.

We investigated the density of STRs detected as a function of GC content across the kingdoms; we observe that by increasing the GC content threshold to 10%, the STR density in Protista drops by 65%, from 1,425.12 to 492.27 STR bps per Mb (**Figure 2c**). The drop in STRs detected as a function of GC content is sharper in eukaryotic organisms with a percentage drop of 41.17%, 44,.42% and 46.11% in Fungi, Animalia and Plantae, respectively in contrast to Archaebacteria, Eubacteria and Viruses (**Figure 2c**; **Supplementary Figure 2**). Moreover, the second large percentage change in STR density occurs with a GC threshold of 60% and 70%. Notably, at 70% GC threshold, STR density in Archaebacteria drops from 21.76 to 15.72 STR bps per Mb (**Figure 3c** **and Supplementary Table 1**). These results can be explained by the higher Adenosine/Thymine (AT) content of eukaryotic genomes, which is reflected in higher AT content in STRs. We conclude that the frequency of short tandem repeats varies significantly across taxa across the domain, kingdom and phyla levels.

**Figure 3:**
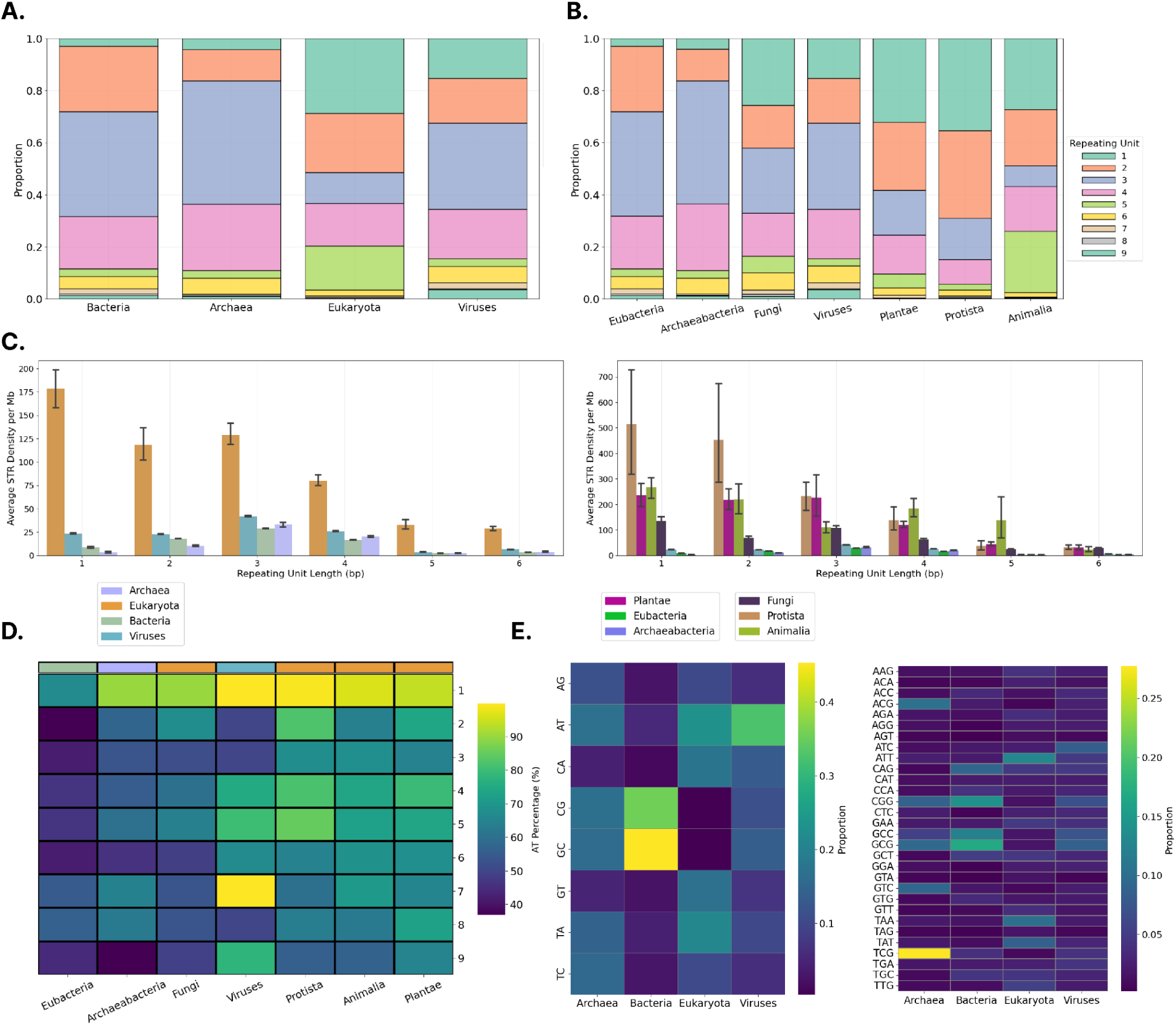
STR density for different repeat unit lengths and nucleotide composition in organisms across the tree of life. **A.** Proportion of total STR occurrences stratified by the repeating unit length for the three domains of life and viruses. **B.** Proportion of total STR occurrences stratified by the repeating unit length per kingdom. **C.** Average density of STRs stratified by the repeating unit length per domain of life and kingdom**. D.** Percentage of AT Content of STRs stratified by the repeating unit length and kingdom. **E.** Proportion of total dinucleotide (left) and trinucleotide (right) STRs partitioned by domain of life, including viruses.

### Significant variation in the abundance of different STR repeat units in different taxa

To investigate potential differences among taxonomic groups based on the most common types of STRs in their genomes, we categorized the STRs according to the length of the repeat unit. We observe that in Eubacteria, Archaebacteria, and Viruses trinucleotide STRs are the most abundant, whereas in Plantae, Protista. Fungi and Animalia mononucleotide STRs are most abundant (**Figure 3a-b**). Of interest is also the fact that in animals the second most prevalent STR type is pentanucleotide STRs (**Figure 3a-b**), which could be driven by telomeric and centromeric regions.

We found that in eukaryotes the most frequent STRs are those with 1bp repeat unit length, whereas in prokaryotic and viral genomes the most frequent are those of 3bp repeat unit length (**Figure 3c; Supplementary Figure 3**). This can be explained by the larger proportion of the genomic space being coding in prokaryotic and viral genomes, in which trinucleotide repeats maintain the reading frame in contrast to mono and dinucleotide repeat units. When partitioning the organismal genomes into the different kingdoms, we find in all eukaryotic kingdoms, mononucleotide STRs to be the most frequent. Notably, in Protista and Animalia the second most frequent is dinucleotide STRs, whereas in Plantae trinucleotide STRs. We also observe a non-negligible density of tetranucleotide and pentanucleotide STR exclusive in animals (**Figure 3c; Supplementary Figure 4**). Using the T2T genomes available, we also observe that certain species display long stretches of STRs that can span hundreds of thousands of bps, which are found in animal and plant species (**Supplementary Table 2**). These findings indicate variation in the frequency of the different STR unit lengths between taxa.

Next, we investigated the AT composition of STRs. We stratified organismal genomes by their kingdoms, and calculated the total percentage of AT content in the STRs of each organism. We observed that mononucleotide repeats are predominantly AT-driven in Viruses, Protista, Animalia and Plantae kingdoms (**Figure 3d**). Notably, the seven bp STRs in Viruses are also highly AT-driven (**Figure 3d**). Upon examining the dinucleotide repeats, the GC and CG tandem repeats were the most frequent dinucleotide repeat type in Bacteria, whereas in eukaryotic organismal genomes, the differences were more sparsely scattered, with TA, AT as well as GT and CA sharing a large proportion of the existing STRs, indicating that there are significant underlying differences between the various kingdoms (**Figure 3e**). Finally, trinucleotide STRs revealed a more equally dispersed distribution, with a notable exception of trinucleotide TCG in Archaea, which are highly enriched (**Figure 3e**). These differences suggested that the analysis could be conducted on the phylum level. Upon further partitioning of the genomes in phyla, we observed that GC-rich STRs are highly specific to a distinct collection of bacterial phyla, whereas other STRs are predominantly AT rich or more uniformly distributed (**Figure 4**). Furthermore, in eukaryotic phyla CA dinucleotide repeats seem to emerge more frequently than the other dinucleotide STR categories, and TA/AT constitute the vast majority of Apicomplexa STRs **Figure 4**). Similar results were observed for trinucleotide STRs, with marked differences being observed both at the domain and phylum levels (**Supplementary Figure 5-6**). Overall, the biophysical properties of STRs are highly divergent across the different domains of life and kingdoms and different STR patterns emerge that are highly specific, indicating that they are involved in different biological mechanisms and molecular processes depending on the organism.

**Figure 4:**
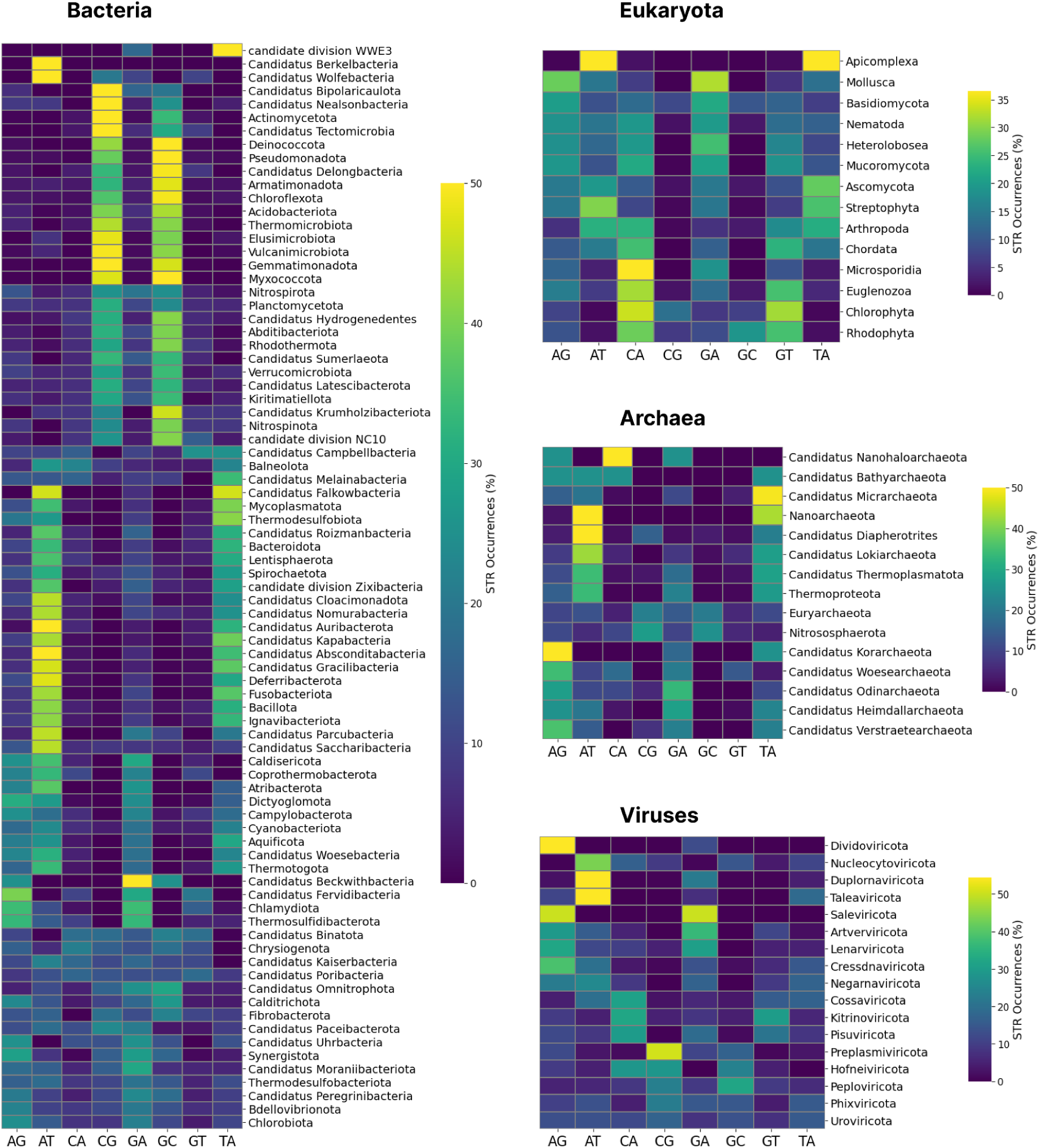
Prevalence of dinucleotide STRs across phyla in the domains of life and viruses. Results shown separately for phyla belonging to each domain of life and in viruses.

### STRs are enriched in eukaryotic and viral, but not in bacterial or archaeal genomes

We next examined if STRs are found more frequently in organismal genomes than expected by chance. To that end, for every organismal genome we generated a matched simulated genome (Jiang et al. 2008), controlling for dinucleotide content, resulting in 117,253 simulated genomes. We quantified the density of different STR categories across taxonomies, at the domain, kingdom and phylum levels. To address the question of prevalence of STRs in a given organismal genome, we compared the resulting STR densities in the real and the shuffled genomes. When examining the three domains of life and viruses, we find that eukaryotes and viruses have on average a 9.79-fold and 1.82-fold enrichment of STRs than expected by chance (Mann-Whitney U test, p-value<0.0001). In contrast, bacteria and archaea show a 0.83-fold and 0,89-fold enrichment, indicating no significant enrichment of STRs in their genomes (Mann-Whitney U test, p-value<0.0001) (**Figure 5a-b**).

**Figure 5:**
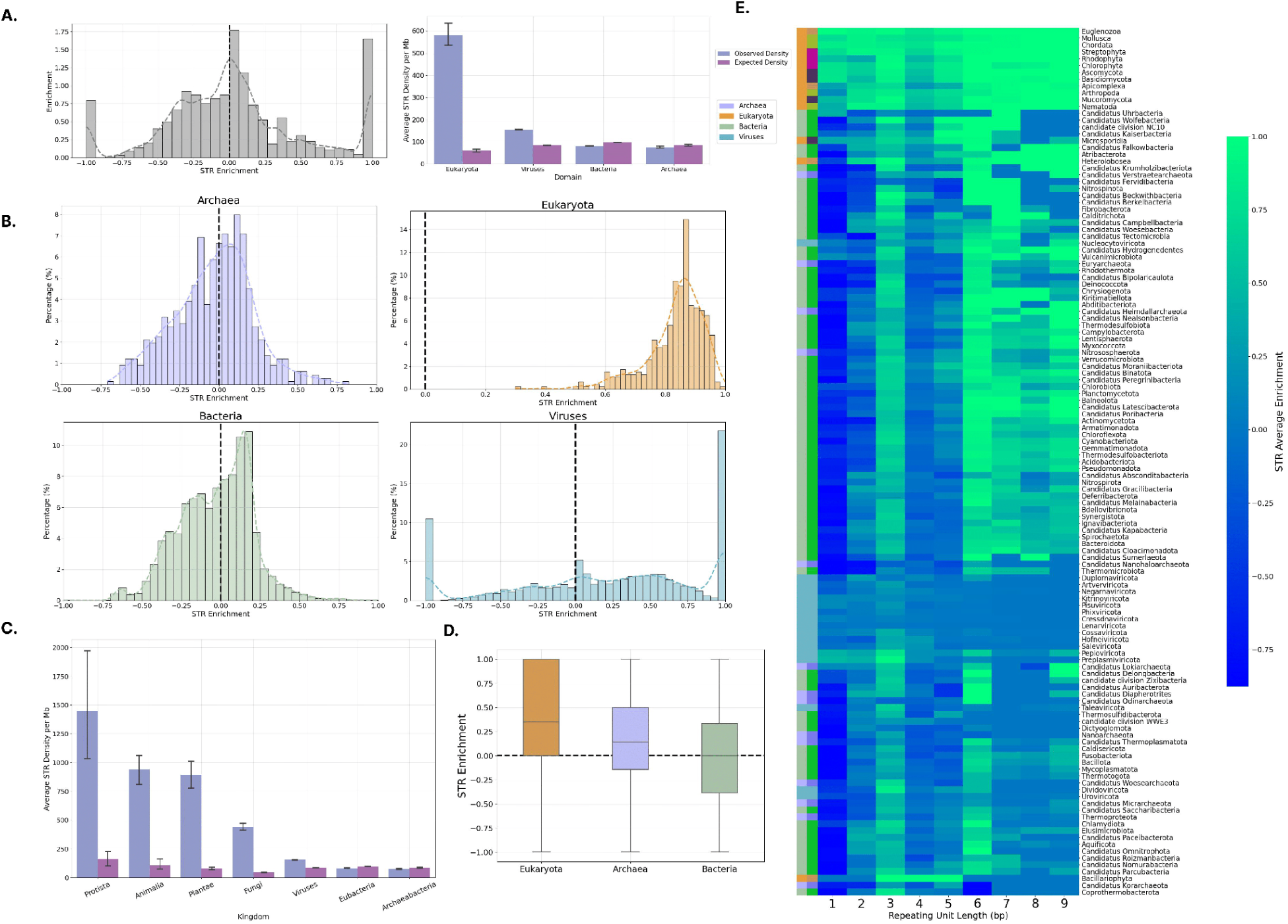
Enrichment of STRs in organismal genomes. **A.** Enrichment of STRs in organismal genomes. **B.** Expected and observed STR density in organismal genomes across the three domains of life and viruses. **C.** STR enrichment in organismal genomes, separated by domain of life. **D.** STR enrichment in viral genomes, separated by their host domain of life. **E.** Expected and observed STR density in organismal genomes across the different kingdoms.

Interestingly, we observe that all eukaryotic species are enriched in STRs, indicating that this is universally the case (**Figure 5b**). Additionally, in bacteria, archaea and viruses there is a significant amount of variance amongst the various organismal genomes (**Figure 5b**). Furthermore, in several cases, viral genomes that were initially found not to contain any STRs, in the simulated genomes contained several STRs and thus obtained a maximum enrichment score of 1.00. Analogously, initially viral genomes that had STRs, after the chromosomal permutations, were rendered empty due to stochastic effects, obtaining the minimum enrichment score of -1.00 (**Figure 5b**). This phenomenon was exclusive to viruses, due to inherently smaller genome sizes of viral genomes, that allowed for such extreme cases to occur. Among eukaryotes, the largest enrichments were observed in Plantae (enrichment=0.83), followed by Fungi (enrichment=0.78), Animalia (enrichment=0.78) and Protists (enrichment=0.74) (**Figure 5c**; **Supplementary Figure 7**). When separating viruses, by their host in those infecting eukaryotes, archaea and bacteria we observe that eukaryotic viruses (219.23 STRs per Mb) have a higher STR density than archaeal (127.86 STRs per Mb) and bacterial (74.36 STRs per Mb) viruses (Mann-Whitney U, p-values<0.0001) (**Figure 5d**). Additionally, archaeal viruses show a significantly higher STR enrichment than bacterial viruses (Mann-Whitney U, p-values<0.0001), the latter of which have the lowest STR enrichment, with the STR density being similar to that of the simulated genomes (**Figure 5d**).

Due to the high variance amongst viral and prokaryotic genomes, we examined how the enrichment of STRs varies in the various phyla, for all repeating unit lengths. We observe a significant amount of enrichment for all eukaryotic phyla, and for all repeat unit lengths. However, for prokaryotic genomes, we notice that one bp repeat unit STRs are highly depleted. Notably, trinucleotide and hexanucleotide tandem repeats are also enriched in most prokaryotic phyla, but not in several viral phyla (**Figure 5e**), which could be explained by the overlapping reading frames of the highly compact viral genomes. This is in contrast with eukaryotes in which one bp repeat unit STRs are highly enriched (**Figure 5e**). Interestingly, there is also a subset of STRs of repeat unit length of six to nine bps that are highly enriched in prokaryotic phyla (**Figure 5e**). These findings indicate that bacterial and archaeal genomes are on average not more repetitive than expected by chance in contrast to eukaryotic and viral genomes.

### STRs are preferentially positioned relative to functional elements in specific taxa

Next, we investigated the distribution of STRs across different genomic elements, including genic, exonic and CDS regions. At the phylum level, for the vast majority of phyla, the STR density is highest when looking on the genome level, rather than any specific genome subcompartment, indicating that STRs are primarily found in intergenic areas (**Figure 6a-b**). This pattern is consistent across unit lengths, with only exceptions being trinucleotide and hexanucleotide STRs, which are more enriched in genic, exonic and CDS regions (**Figure 6a-c**). For Chordata, Streptophyta, Chlorophyta and Apicomplexa we observe high STR densities in genic and exonic regions. In particular, for Chlorophyta and Apicomplexa, we observe high STR densities in CDS regions, besides genic and exonic (**Figure 6c**). In prokaryotic genomes we observe a general depletion of STRs, with several notable exceptions (**Figure 6c**). For instance, the bacterial phyla Deinococcota and Bacteroidota, display a high genome-wide STR density, and similarly high genic/CDS density of STRs. We also observe that trinucleotide unit STRs are enriched across these compartments, which is explained by the selection against disrupting reading frames (Coenye and Vandamme 2005; Lin and Kussell 2012; Mrázek, Guo, and Shah 2007).

**Figure 6:**
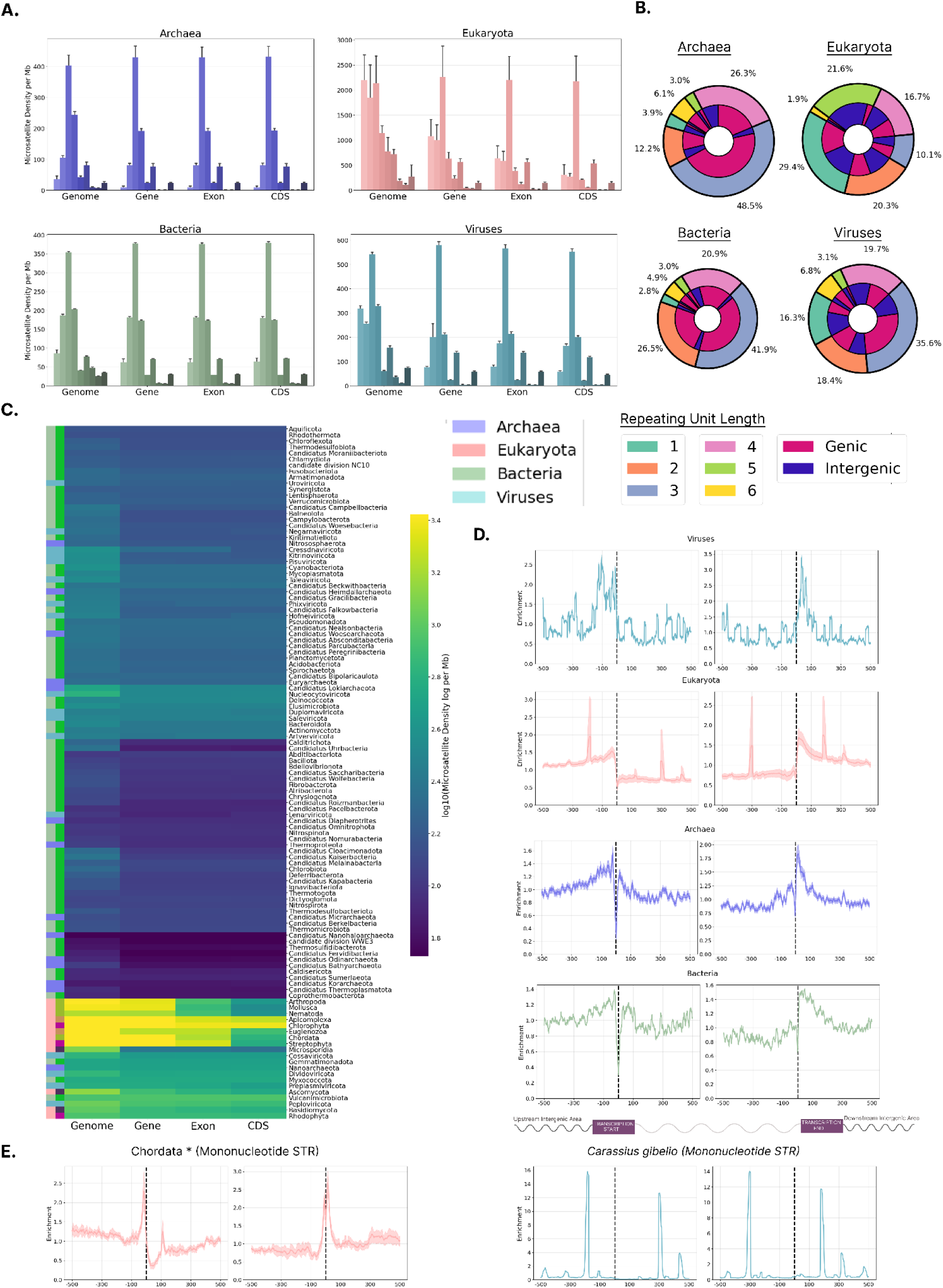
The topography of STRs relative to genomic subcompartments and transcription start and transcription end sites, across taxa. **A.** STR density for the three domains of life and viruses at the genome, genic, exonic and coding regions. **B.** Pie charts showing the percentage of STRs separated by STR unit length and proportion being genic and intergenic for the three domains of life and viruses. **C.** STR density of organisms belonging to different phyla at the genome, genic, exonic and coding regions. **D.** STR distribution across the three domains of life and viruses, relative to the transcription start and end sites. **E.** Mononucleotide STR distribution in chordate (excluding *Carassius gibelio*) and in *Carassius gibelio*. Error bars represent the 2.5% lowest and 97.5% highest percentile from Monte-Carlo simulations with replacement (N=1,000).

### Enrichment of STRs relative to transcription start and end sites

Provided that the distribution of STRs was found to be uneven across genomic subcompartments, we investigated the distribution of STRs relative to functional genomic elements including Transcription Start Sites (TSSs) and Transcription End Sites (TESs). Our hypothesis was that if STRs have functional roles they will be enriched relative to key genomic loci. We find that viruses are enriched upstream of the TSS and downstream of the TES. In bacteria and archaea weaker effects were observed, with weak enrichments found upstream and downstream of genic regions (**Figure 6d**). Eukaryotes showed strong and highly localized enrichment at roughly 150bps upstream from the TSS, 300bps downstream from the TSS, and similarly roughly 300bps upstream of the TES and 150bps downstream of the TES (**Figure 6d**).

Nonetheless, around the modes, the confidence intervals were highly volatile, suggesting that the observed signal originating from a certain subset of species and putatively a STR of specific length. This led us to separate the eukaryotic phyla, partition the distributions across the various repeating unit lengths of STRs and examine separately the STR enrichment relative to the TSS and TES, finding significant differences between them. Examination of the density of mononucleotide STRs amongst Chordata revealed that this sharp enrichment was stemming from mononucleotide STRs in *Carassius gibelio*, whereas the other members of the Chordata displayed an enrichment at the TSS and TES (**Figure 6e**). Subsequent examination of the remaining eukaryotic phyla, showed that Ascomycota and Apicomplexa have polyA and polyT mononucleotide STRs enriched directly at the TSS and TES. However, Streptophyta mononucleotide repeat distribution was comparable to Chordata, but the polyA repeats were, instead, replaced with polyT repeats (**Supplementary figure 9**).

### Large differences in STR frequency across and within human chromosomes

Using the T2T complete human genome (Nurk et al. 2022), we examined the distribution, frequency and diversity of STRs across the complete chromosomes including low complexity regions such as pericentromeric, centromeric and telomeric loci, which have been traditionally difficult to accurately sequence and study (Logsdon et al. 2023).

In total we report 3,612,909 STRs in the T2T human genome. We investigated the STR composition and density across and within each chromosome. To examine the relationship between STRs and centromeres, each chromosome was annotated by the relevant pericentromeric and centromeric positions. We report marked differences in STR distribution patterns between and within chromosomes. Strong STR enrichments are observed in chromosomes 21 and 22, in the short chromosome arms. Particularly striking is the cluster of STRs in the short arm of chromosome 21 (21p) at Ribosomal DNA (rDNA) genes. In contrast, chromosomes 1, 2, 3, 4, 5, 7, 10, 16, 17 and 20 show sharp STR peaks in the vicinity of the centromeres and in pericentromeric regions (**Figure 7**; **Supplementary Figure 12**). For chromosomes 6, 8, 11, 12, 18 and 19 we do not find strong STR enrichments at any part of the chromosome.

**Figure 7:**
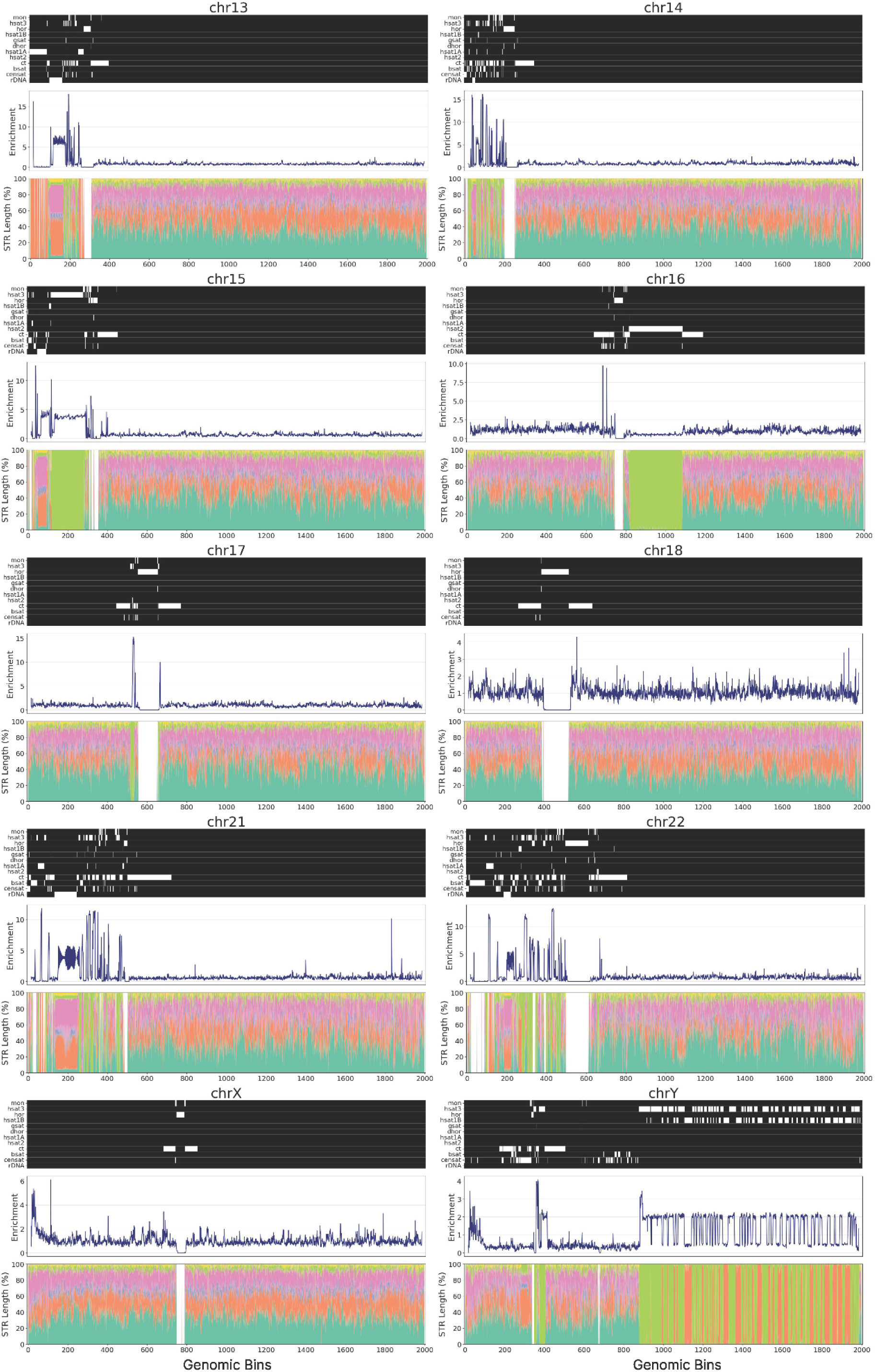
Characterization of STRs across chromosomes in the T2T reference human genome. Schematics show the distribution of STRs across different human chromosomes. The heatmap shows the different types of pericentromeric and centromeric repeats, with white color representing presence of the repeat in that genomic region. Line plots show the STR enrichment at each genomic bin for a chromosome. Stacked barplots show the results for different STR unit lengths. Repeats include inactive αSat HOR (hor), divergent αSat HOR (dhor), monomeric αSat (mon), classical human satellite 1A (hsat1A), classical human satellite 1B (hsat1B), classical human satellite 2 (hsat2), classical human satellite 3 (hsat3), beta satellite (bsat), gamma satellite (gsat), other centromeric satellites (censat) and centromeric transition regions (ct).

In the acrocentric chromosomes 13, 14 and 15, the HSat3 arrays span the centromere, displaying strong enrichment on the short arm. Interestingly, in the aforementioned chromosomes, the rDNA array genomic region, which is proximal to the centromere, shows a particularly strong STR enrichment, with the exception that its corresponding STR composition is highly diverse; a phenomenon which is in contrast to other highly repetitive STR-enriched regions where the STRs are biased towards a specific repeat unit length. Interestingly, chromosome 9 is an outlier, with the whole centromere being composed of five bp unit STR repeats, with this region displaying the highest enrichment in the chromosome. The Y chromosome displays a strong STR enrichment in over half of its length, which are composed primarily of dinucleotide and tetranucleotide STRs (**Figure 7**; **Supplementary Figure 12**). Amplicons are highly prevalent in mammalian Y chromosomes (Skaletsky et al. 2003) and account for the repeated pattern of STR enrichment with intervals over long genomic distances (**Figure 7**; **Supplementary Figure 12**). These results indicate the highly differing STR profile of the different human chromosomes, while acrocentric chromosomes show a distinct and consistent pattern of high STR density in the shortest arm.

Next, we investigated the STR density across human genomic subcompartments, including the different types of satellite array regions that are part of the pericentric and centromeric repeats. We find that STRs are most enriched in telomeric regions (**Figure 8a**), which is consistent with them being composed of STRs (Rosenberg et al. 1997). These are followed by classical human satellite 3 (hsat3) repeats and rDNA loci (**Figure 8a**). Interestingly, silencers and inactive and divergent alpha satellite higher order repeats show the strongest depletion for STRs (**Figure 8b**). When examining each repeat length separately we find that hexanucleotide STRs are most prevalent in telomeres, which is expected, and STRs of five bp repeat unit length are most prevalent in classical human satellite 2 (hsat2), hsat3 repeats. In contrast, dinucleotide STRs are enriched at hsat1A and hsat1B sites and mononucleotide STRs in gamma satellites (gsat) (**Figure 8b**).

**Figure 8:**
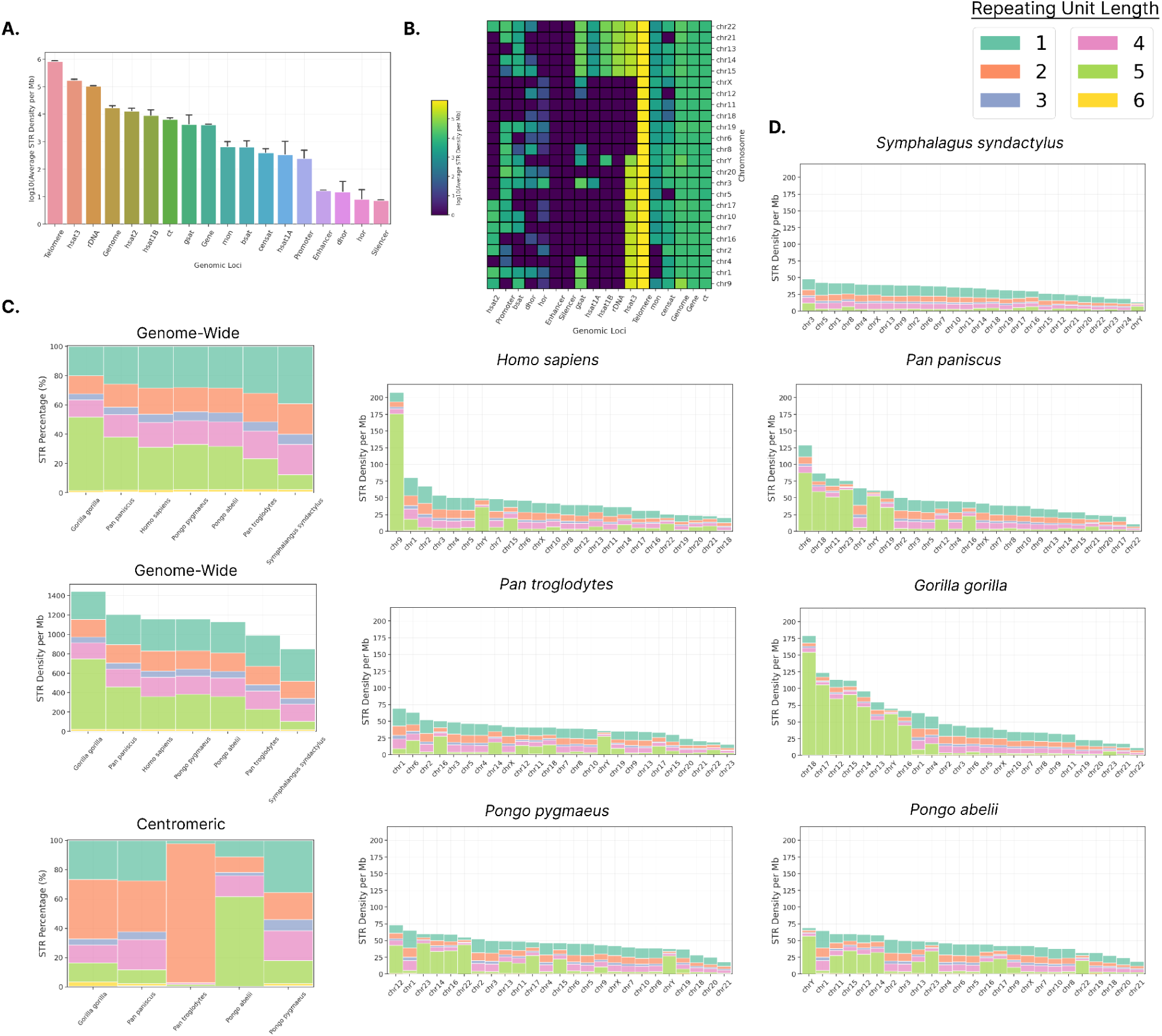
Distribution of STRs in Telomere-to-Telomere primate genomes. **A.** STR Density across human genome sub-compartments including centromeric repeats. **B.** STR Density across human genome sub-compartments including centromeric repeats split by chromosome. **C.** Percentage and density of different lengths of STRs genome-wide and in centromeres. **D.** STR density for STRs of different unit length across primate species, in each of their chromosomes. Repeats include inactive αSat HOR (hor), divergent αSat HOR (dhor), monomeric αSat (mon), classical human satellite 1A (hsat1A), classical human satellite 1B (hsat1B), classical human satellite 2 (hsat2), classical human satellite 3 (hsat3), beta satellite (bsat), gamma satellite (gsat), other centromeric satellites (censat) and centromeric transition regions (ct).

### T2T primate genomes reveal rapid and dynamic evolution of STRs in the primate lineage

Next, we used the genomes from the T2T-Primates consortium (Hoyt et al. 2022), and investigated the distribution of STRs in six non-human primate genomes including *Gorilla gorilla* (gorilla), *Pan paniscus* (bonobo), *Pan troglodytes* (chimpanzee), *Pongo abelii* (Sumatran orangutan), *Pongo pygmaeus* (Bornean orangutan) and *Symphalangus syndactylus* (Siamang gibbon). Among primates we find that *Gorilla gorilla* shows the highest STR genome-wide density, whereas *Symphalangus syndactylus* has the lowest genome-wide STR density (**Figure 8b**).

Centromeres have arisen during eukaryotic evolution that are some of the most fast evolving regions of the genome and therefore were particularly interested to examine the STR diversity and distribution in centromeric and pericentromeric repeats across the primate species. We also find that in multiple primate species, including humans, gorilla and bonobo, pentanucleotide STRs are the most abundant type of STR (**Figure 8b**). In repeats found in centromeric and pericentromeric regions dinucleotide STRs are the most abundant followed by pentanucleotide and dinucleotide STRs depending on the species and the genomic compartment (**Figure 8b**; **Supplementary Figure 13**).

Recent work has indicated that the sex chromosomes are highly repetitive and has examined the presence of long palindromes, endogenous repeat elements and satellites (Miga et al. 2020; Rhie et al. 2023; Makova et al. 2023). However, the contribution of STR repeat types has not been examined. Here we focused on the distribution of STRs throughout the sex chromosomes in the primate lineage. Interestingly, the STR profile of chromosome Y is remarkably different between the human and non-human primate genomes (**Figure 7**-**8**, **Supplementary Figure 12, Supplementary Figure 14**). In contrast, the STR content of chromosome X is markedly similar between these genomes (**Figure 8**, **Supplementary Figure 14-15**).

## Discussion

In this study, we conducted a comprehensive analysis of STR variations across 117,253 organismal genomes from a broad range of organisms spanning all major taxonomic groups. Our findings reveal that STRs are predominantly enriched in eukaryotic organisms, with the highest STR density observed in *Plasmodium falciparum*. We found an extremely high variance between domains, with eukaryotic organisms having the highest STR density and archaea the lowest. Interestingly, similar variation in STR density was identified upon partitioning the viruses with respect to the host domain. Viruses of eukaryotic hosts had a disproportionately higher STR density than the corresponding viruses of bacterial and archaeal hosts. *O*n the kingdom level, Protista had the highest average STR density followed by Animalia. Upon further partitioning on the phylum level, Euglenozoa had the highest average STR density followed by several other eukaryotic phyla. The bacterial and archaeal genomes have fewer intergenic regions and most prokaryotic phyla showed low STR densities, with some notable exceptions, in accordance with previous literature (Field and Wills 1998; Schlötterer et al. 2006; Mrázek, Guo, and Shah 2007).

STRs are drivers of genome evolution and adaptation (Verbiest et al. 2023; Zhou, Aertsen, and Michiels 2014), are highly dynamic elements and can expand or contract primarily due to strand-slippage replication events. Therefore, slippage events can result in the addition or deletion of repeat units, leading to insertions or deletions. These mutations can have various effects, ranging from benign to harmful, depending on their location and contribute to human genetic disorders, especially those characterized by repeat expansions, such as Huntington’s disease and muscular dystrophy (Tanudisastro et al. 2024). Thus, understanding how different organisms cope with the genomic instability is of importance. Through genome simulation experiments, we discovered that archaea and bacteria typically exhibit fewer STRs than expected, while eukaryotes possess a roughly ten-fold enrichment. When separating viruses by their host domain, eukaryotic viruses show higher STR enrichment than archaeal and bacterial viruses. We also observed that STRs tend to be located near functional genomic elements in specific taxa. These results highlight substantial variations in the frequency and distribution of STRs across the tree of life. It should be noted that there were considerable differences in enrichment of STRs across the various phyla, potentially indicating the STRs have evolved to assist different functionalities and biological mechanisms in different phyla or that they are more likely to expand in certain phyla than others. These results are consistent with the current literature, where STRs have an uneven distribution in prokaryotic genomes (Verbiest et al. 2023).

The frequency of STRs varies significantly for different repeat unit lengths and is influenced by the organism’s taxonomic classification. Several STR motifs were far more prevalent when focusing on a specific domain. For instance, dinucleotide STRs in bacteria, most commonly emerge with the consensus GC or CG. In general, STRs in prokaryotes were GC-enriched in contrast to other eukaryotic or viral species, due to the genome GC content differences. We also examined if STRs are preferentially located at certain genomic compartments such as genic, exonic and CDS regions. By partitioning STRs into unique sets of one to nine bps unit length, we observed that across all prokaryotic organisms, trinucleotide repeats are the most prevalent in CDS regions, which is expected for maintaining the reading frame in coding regions. In several eukaryotic and viral species, the genic density of STRs was higher than the corresponding CDS density, leading us to the conclusion that STRs have a higher density in intronic and intergenic rather than exonic areas. Furthermore, mononucleotide repeats constituted 29.4% of total STRs in eukaryotic species, followed by 21.6% of pentanucleotide STRs and 20.4% of dinucleotide STRs. Dinucleotide and pentanucleotide STRs were concentrated in non-genic areas, as indicated in the human genome and non-human primate genomes, in which they were highly enriched in centromeric and pericentromeric regions.

The distribution of STRs relative to TSSs and TESs was also highly variable depending on the taxon. Most interestingly, in eukaryotic species, an enrichment relative to the TSS and TES led us to segment further into phyla and individual species in our analysis. To our surprise, the signal originated from a goldfish, which had an enrichment of mononucleotide repeats roughly 150bp upstream from the TSS. Various protists and fungi phyla displayed enrichment of STRs relative to the TSS, with mononucleotide and dinucleotide repeat density reaching the highest enrichment just before the TSS and just after the TES. This is in contrast to other eukaryotic genomes originating from animal or plant phyla, such as Streptophyta or Chordata, where the transcriptional regulatory roles of STRs were far more diverse with several positions relative to transcription start and end sites being enriched. In Chordata, mononucleotide repeats relative to the TSS mostly consist of polyA tails, whereas in Streptophyta polyT tails. Evolutionary Streptophyta seem to have diverged from other eukaryotic phyla, by replacing most A-rich polyadenylation signals with U-rich motifs (Wodniok et al. 2007), which could explain the observed differences (Wodniok et al. 2007). In bacteria and archaea, STRs were preferentially positioned downstream of the TESs. These results suggest that STRs repeats play a crucial role in various underlying biological mechanisms but their regulatory roles vary greatly across the different phylum, kingdom and domain taxonomic levels.

Finally, we use the recently completed T2T genomes of various primate species, including the latest assembly of the human genome, to discover that STRs are highly abundant and variable between primate species, especially in peri/centromeric regions. Our findings indicate that STRs are highly dynamic, fast-evolving elements facilitating organismal evolution. The vast differences observed at Y chromosomes amongst the Great Apes and Homo Sapiens are suggestive of its intrinsic plastic nature. The Y chromosome in *Homo Sapiens* is covered by HSat1-3 arrays which account for most of its part. On Y chromosome satellite arrays HSat2-3 being enriched in pentanucleotide repeats whereas hsat1A and hsat1B are enriched in dinucleotide repeats is a common feature amongst primates. The observed differences on the Y chromosome could be attributed to the tendency of heterochromatin to be highly mutagenic with observed differences even amongst homologous chromosomes. Such differences are not observed in the X chromosomes, which STR are more uniformly distributed and far more diverse for the various repeating unit lengths. The high observed variance in Y chromosomes amongst primates could be attributed to the highly mutagenic nature of heterochromatin. In such regions, due to the highly repetitive nature of STR-rich loci, double strand breaks repaired by homologous recombination using the sister chromatid is more likely to cause mutations, which could explain the increased divergence in sequence composition.

As more T2T genomes become available, both for more individuals in a given species and for more diverse species, future studies will be able to further reveal the contribution of STRs in driving eukaryotic evolution and adaptation. To conclude, this study analyzed the distribution and topography of STRs across 117,253 genomes from various organisms, finding that STRs are mostly enriched in eukaryotes, with viruses infecting eukaryotic hosts also showing higher STR densities.The highest STR densities were observed in the Protista kingdom and Euglenozoa phylum, while bacterial and archaeal genomes showed low STR densities. Simulation experiments showed a relative depletion of STRs from bacterial and archaeal genomes.

## Materials and Methods

### Data retrieval and parsing

Complete genomes were downloaded from the GenBank and RefSeq databases (O’Leary et al. 2016; Benson et al. 2013). A total of 117,253 complete organismal genomes were analyzed and integrated in the database. For each genome, the associated files for RNA and coding regions as well as the GFF gene annotation file were downloaded.

We also downloaded T2T genomes for the following primates through the T2T-Primates consortium (Hoyt et al. 2022), *Gorilla gorilla, Pan paniscus*, *Pan troglodytes*, *Pongo abelii*, *Pongo pygmaeus* and *Symphalangus syndactylus* from https://github.com/marbl/Primates. Associated files including genome annotations were downloaded from https://github.com/marbl/t2t-browser.

### Identification of STRs

STRs were detected using RPTRF, a perfect tandem repeat finder tool (Behboudi, Nouri-Baygi, and Naghibzadeh 2023). We used RPTRF to extract the raw perfect STRs from each complete genome available in NCBI Genbank and RefSeq databases with parameters maximum motif size, M=50,000, and minimum length, t=1. The pipeline was split across multiple nodes on the High-performance computing system, each node processing a distinct set of complete genomes. The C++ script processed each chromosome in parallel, passing the derived sequences into a custom Python script. The Python script was created to validate, parse and store all STRs and derive the STRs by bounding the consensus sequence length between one and nine base pairs. Furthermore, consensus sequences containing N or other characters not originating from the nucleotide alphabet {’A’, ’G’, ’C’, ’T’} were filtered out from the extracted STR dataset. Finally, another custom python script was integrated into the existing pipeline to extract the perfect STRs of mononucleotide repeats, and then concatenated the two outputs into a single dataset which was saved in a parquet-snappy compressed format.

### Identification of STRs in simulated organismal genomes

To estimate the expected amount of STRs we used the uShuffle package (Jiang et al. 2008). For each available organismal genome we shuffled all the chromosomes, while preserving the initial dinucleotide frequencies, resulting in simulated control genomes. In each of the simulated genomes we extracted the STRs using the aforementioned RPTRF software. Then, we compared the STR density per Mb across the shuffled genomes and the NCBI organismal genomes, by addressing the enrichment of the observed versus the expected tandem repeat counts. For that reason, we utilized the following formula:

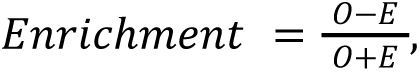

where *O* = *Observed* and *E* = *Expected* STR in a given organismal genome. Given the definition above, the enrichment function ranges from -1 to 1, indicating absolute depletion and absolute enrichment, respectively. In cases where the genome initially did not contain any STRs, and its corresponding shuffled genome was also empty, it was natural to assume that there is neither depletion nor enrichment, and thus, in such cases where *O* = *E* = 0, we set *Enrichment* = 0.

### Estimation of STR density across genomes and genomic subcompartments

STR density in a single organismal genome was determined by dividing the total number of detected STR occurrences by the total base pairs of its genome size, multiplied by 1,000,000. We calculated the average STR density across the genomes within each taxonomic group. Additionally, STR density across different genomic subcompartments was assessed by dividing the total overlap number of base pairs of STRs within each subcompartment by the overall length in base-pairs of these subcompartments. Coordinates for subcompartments were sourced from the relevant GFF files, and any overlapping annotations within a subcompartment were not double-counted. The average STR density was also computed across species within the same taxonomic group, either throughout the entire genome or within specific genomic subcompartments. Species not associated with a GFF file were omitted from this analysis. Furthermore, certain viral species that did not contain any relevant genomic compartment were also skipped during the density extraction.

For each chromosome for the T2T genomes, we generated N=2,000 genomic bins of equal length. Each STR start and end coordinate was assigned at a particular bin from 1 to 2,000, according to the following formula:

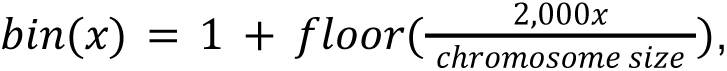

where x is the start or end coordinate of the STR. Subsequently, for each bin the total number of STR occurrences was estimated by summing the total distinct start or end coordinates within a particular bin. In cases where the start and the end of the STR were assigned a different bin, the STR was counted in all of the intermediate bins. We calculated the number of bps covered by STRs, for each STR unit length.

### Estimation of PWM across genomes

The PWM for STR motifs was extracted at a generated window of 500bp upstream and downstream from TSSs or TESs. The process involved a custom python script which utilized bedtools to extract the intersections of tandem repeats with genic compartments derived from GFF files. At each intersection, the tandem repeat sequence was saved as a separate column, along with the corresponding gene strand location. Finally, the script constructed a numpy vector array (1,1001) which stored the occurrences for each nucleotide of tandem repeats relative to the TSSs and TESs coordinates at a particular position, ensuring the motif is translated to the same strand as to where the genic region lies. This process was repeated for repeating unit lengths 1 to 9, for all the available organismal genomes in the NCBI database associated with a GFF file. For the creation of logoplots we used the Logomaker Python package in combination with Matplotlib (Tareen and Kinney 2020). The (4, 1001)-matrix containing the probability of a nucleotide occurring at each position in the expanded window, was calculated by using the Bayes estimator:

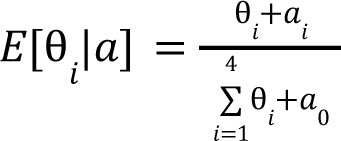

of a Dirichlet prior, where, α = (α_1_, α_2_, α_3_, α_4_) is the pseudocount vector, 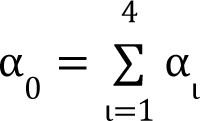 and θ = (θ_1_,θ_2_,θ_3_,θ_4_) is the probability vector of nucleotide from the nucleotide alphabet Π = {‘*A*’, ‘*C*’, ‘*T*’, ‘*G*’,} occuring at a particular position. In our analysis, we used the Dirichlet prior α = (1, 1, 1, 1) as the prior distribution for the probability of each nucleotide occurring at each position.

### Estimation of STR density relative to TSSs and TESs

To investigate the relationship between STR sites and TSSs or TESs, we generated local windows around TSSs/TESs and measured the distribution of STR base pairs across the window. The enrichment was calculated as the number of occurrences at a position over the mean number of occurrences across the window. Confidence intervals were calculated as the 2.5% lowest and 97.5% highest percentile from Monte-Carlo simulations with replacement (N=1,000), in which we randomly picked an equal number of species from the domain, kingdom or phylum that was studied.

## Code Availability

The GitHub code and all the related material is provided at: https://github.com/Georgakopoulos-Soares-lab/treeoflifeSTR

## Author Contributions

N.C., and I.G.S. conceived the study. I.G.S. supervised the study and provided the resources. N.C. generated the figures and tables. N.C., and I.G.S. wrote the manuscript.

## Supplementary Material

**Supplementary Figure 1:**
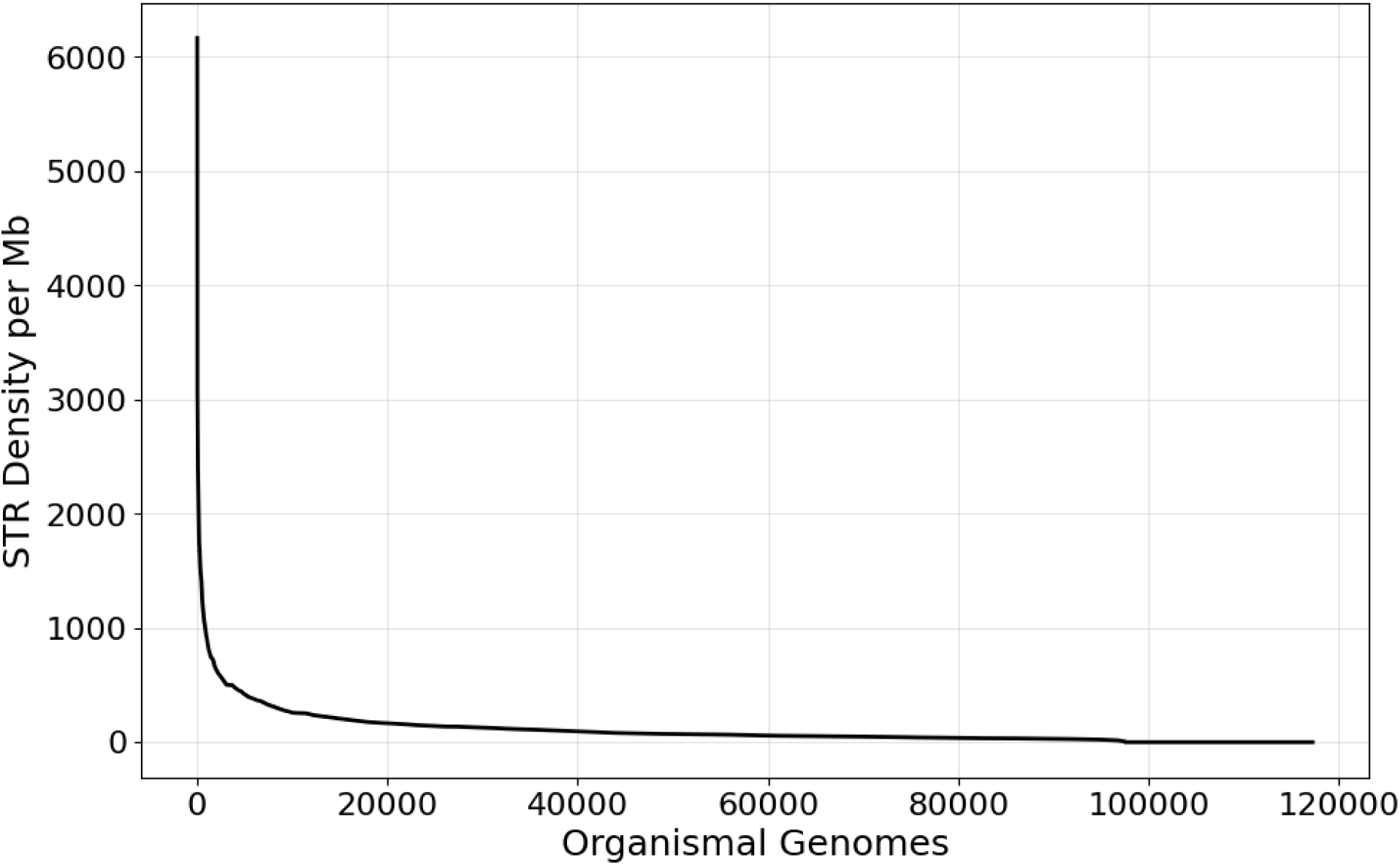
STR density per Kb per organismal genome.

**Supplementary Figure 2:**
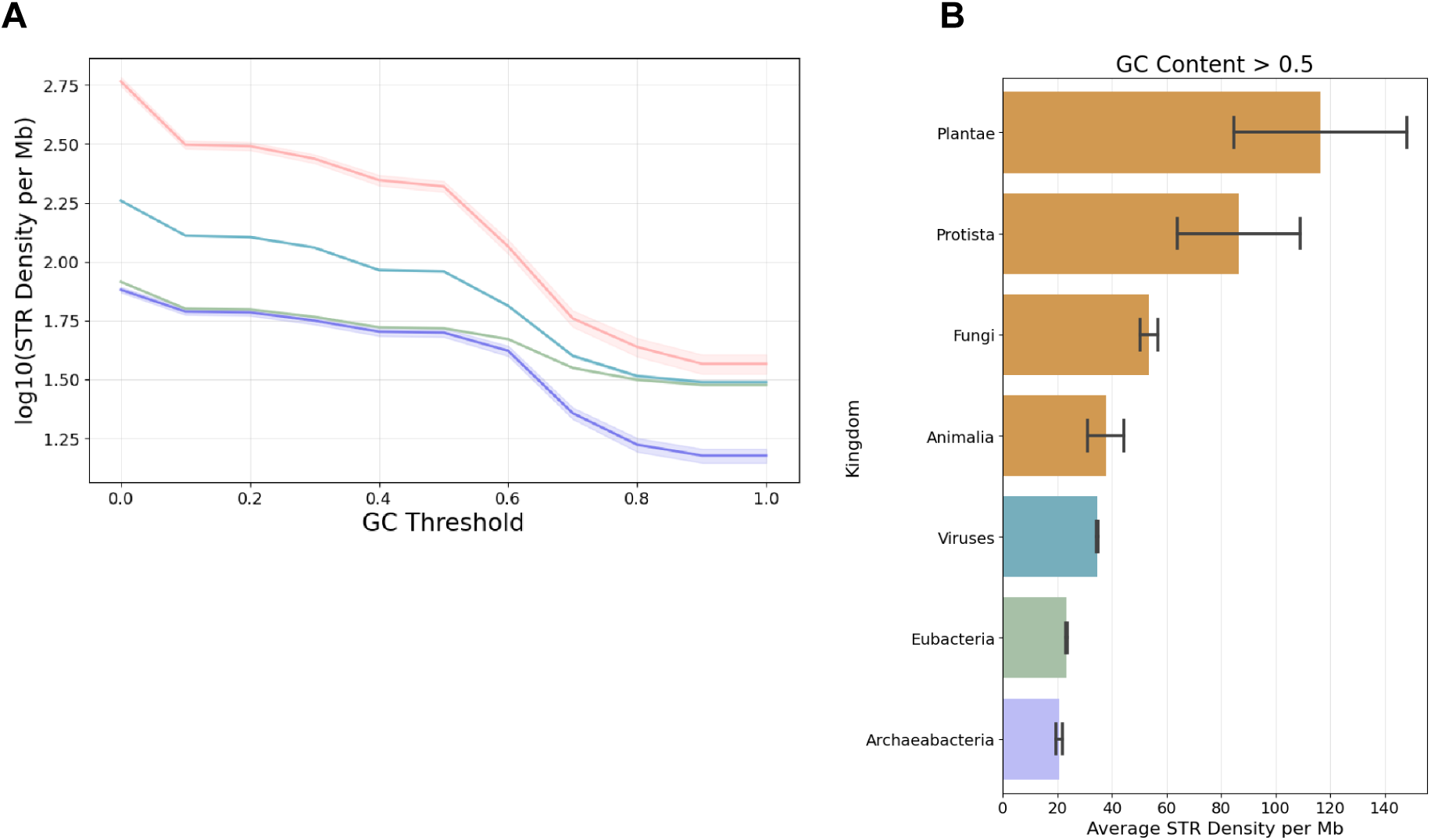
STR density for various GC Content thresholds. Results shown **A.** for the three domains of life and viruses and **B.** for kingdoms. Coloring in B. is done based on the domain of life.

**Supplementary Figure 3:**
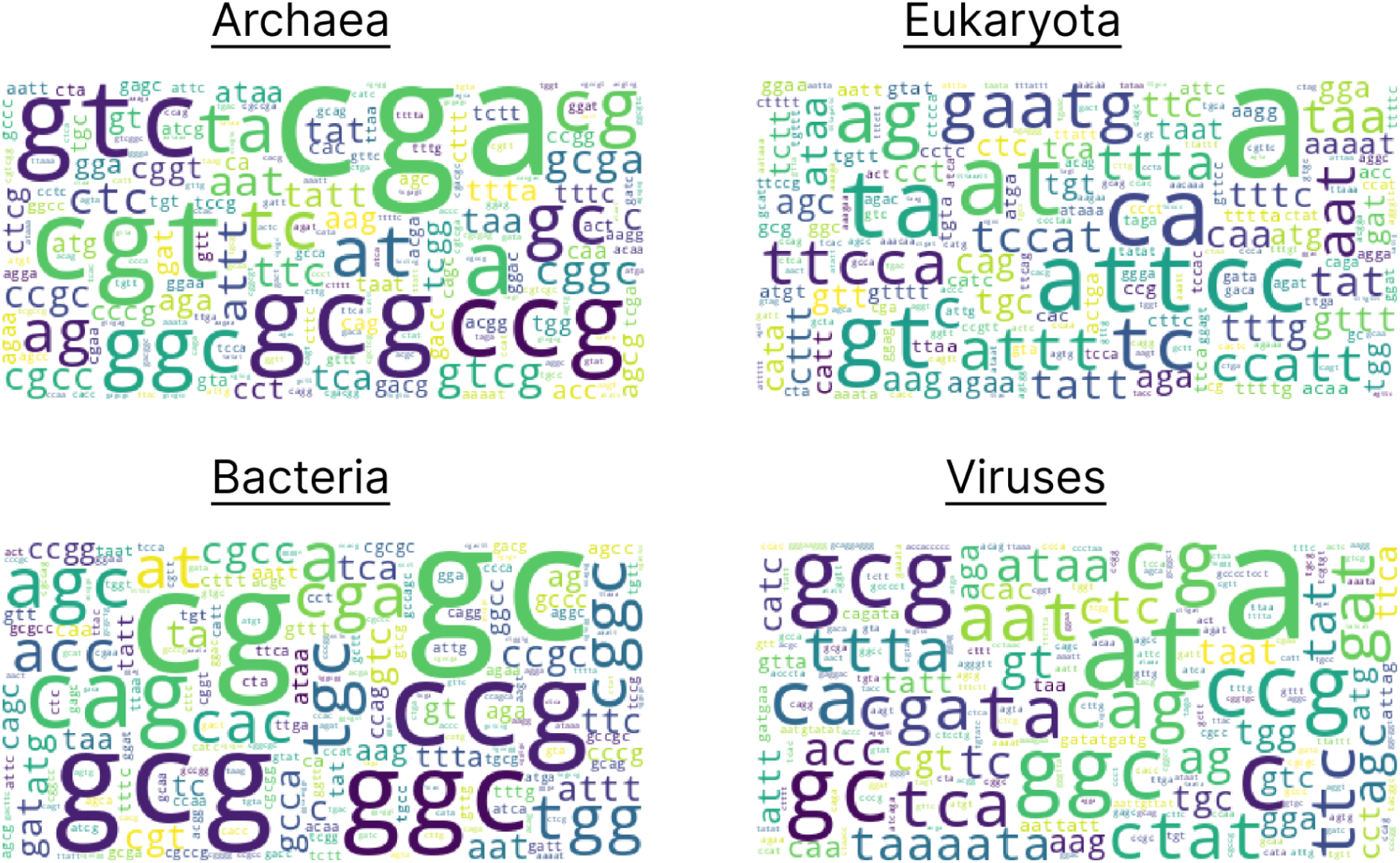
Frequency of most frequent STR motifs across the three domains of life and viruses.

**Supplementary Figure 4:**
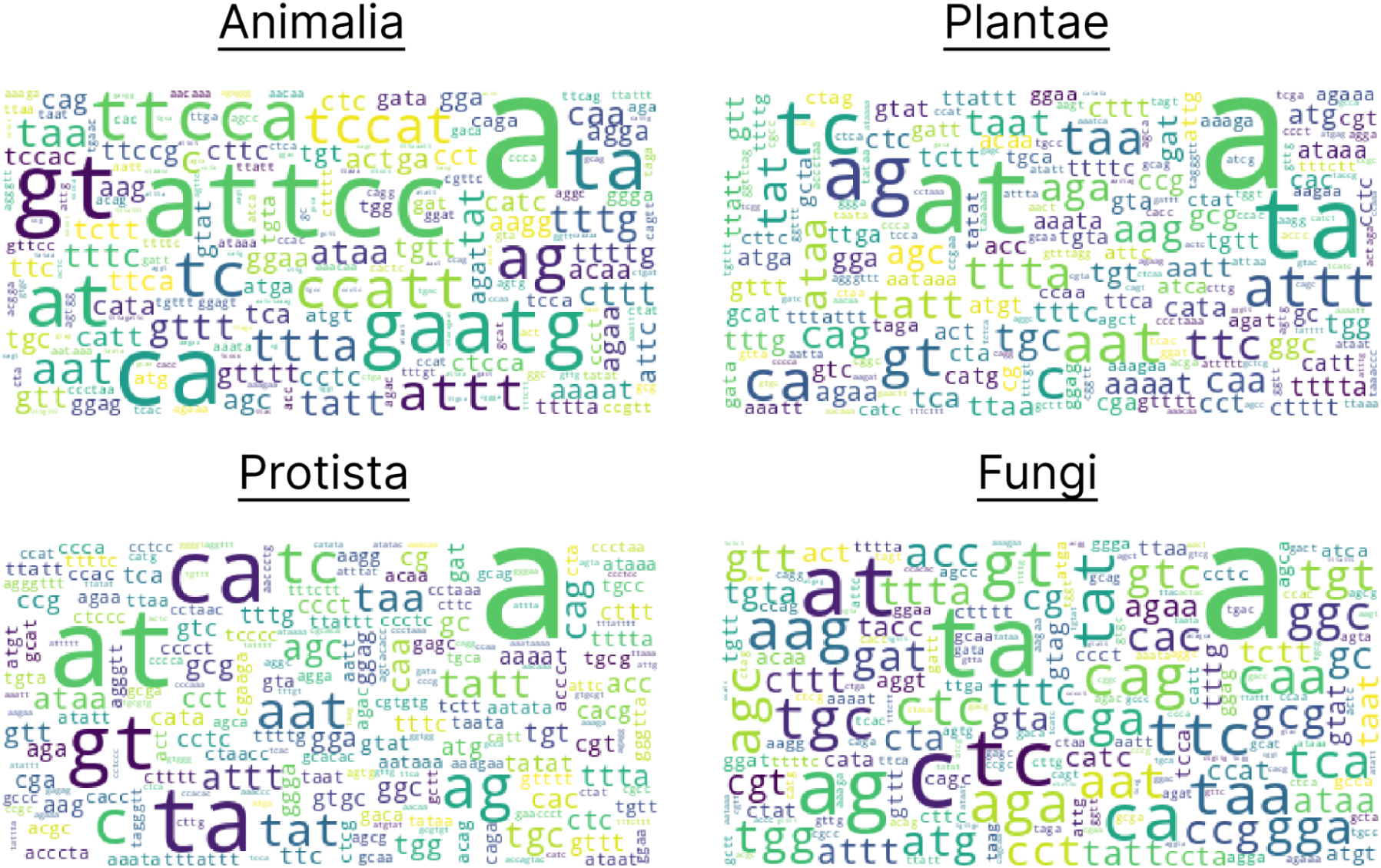
Frequency of most frequent STR motifs across eukaryotic kingdoms.

**Supplementary Figure 5:**
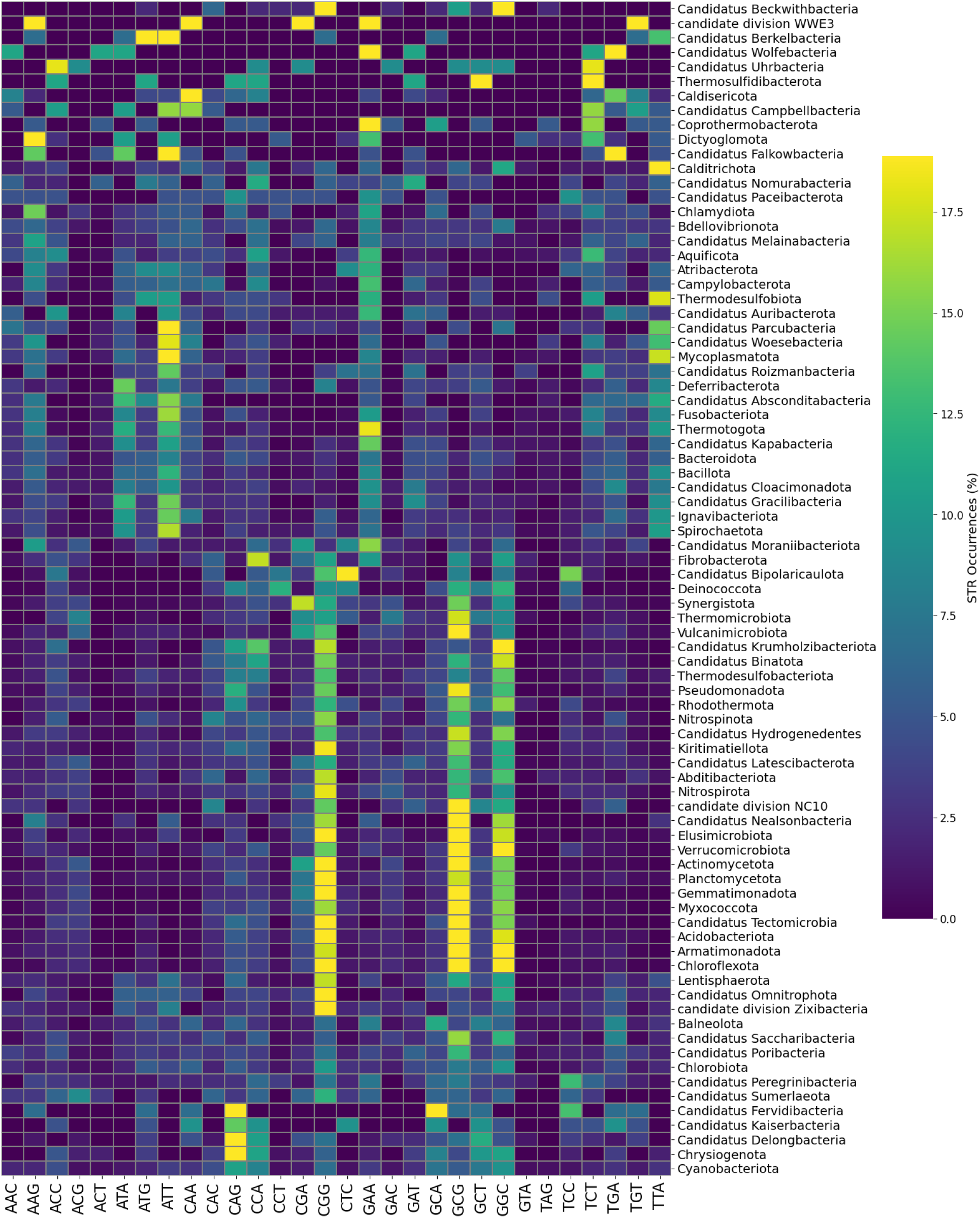
Prevalence of trinucleotide STRs across bacterial phyla.

**Supplementary Figure 6:**
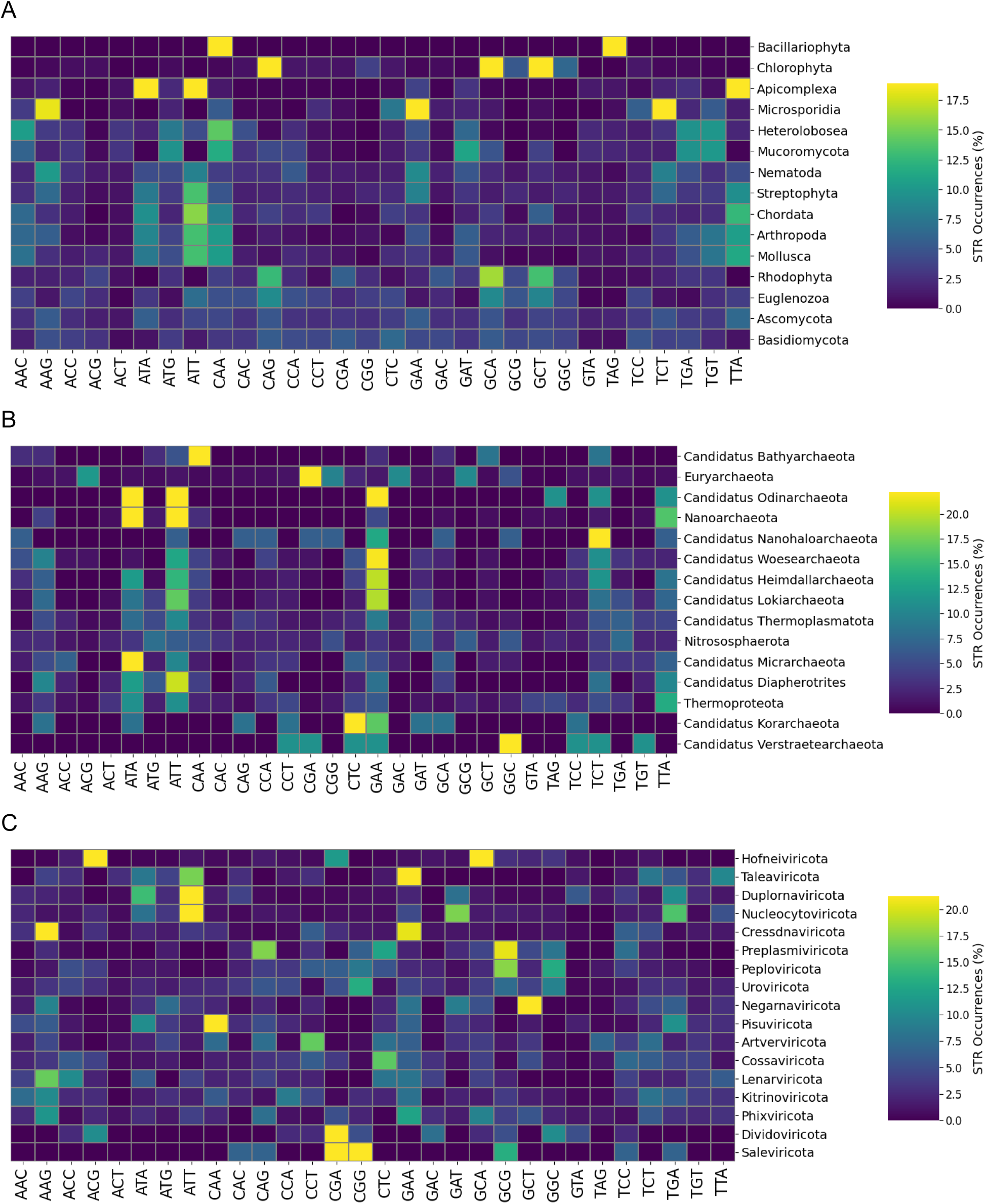
Prevalence of trinucleotide STRs across A. eukaryotic, B. archaeal and C. viral phyla.

**Supplementary Figure 7:**
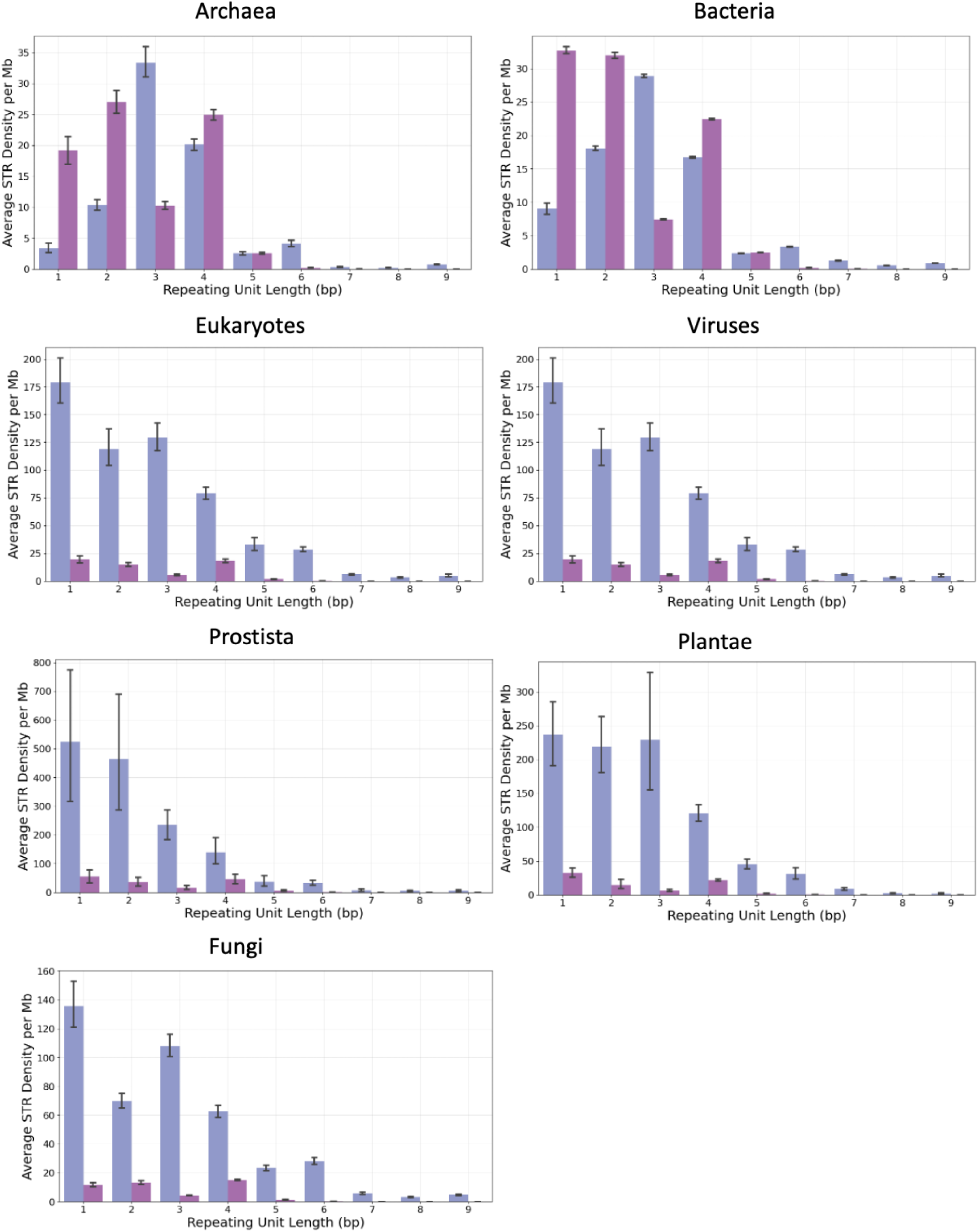
Observed versus expected STR density across different taxa. Results shown for repeat unit lengths of one to nine bps. Blue shows STR density in real genomes and purple indicates STR density in simulated genomes.

**Supplementary Figure 8:**
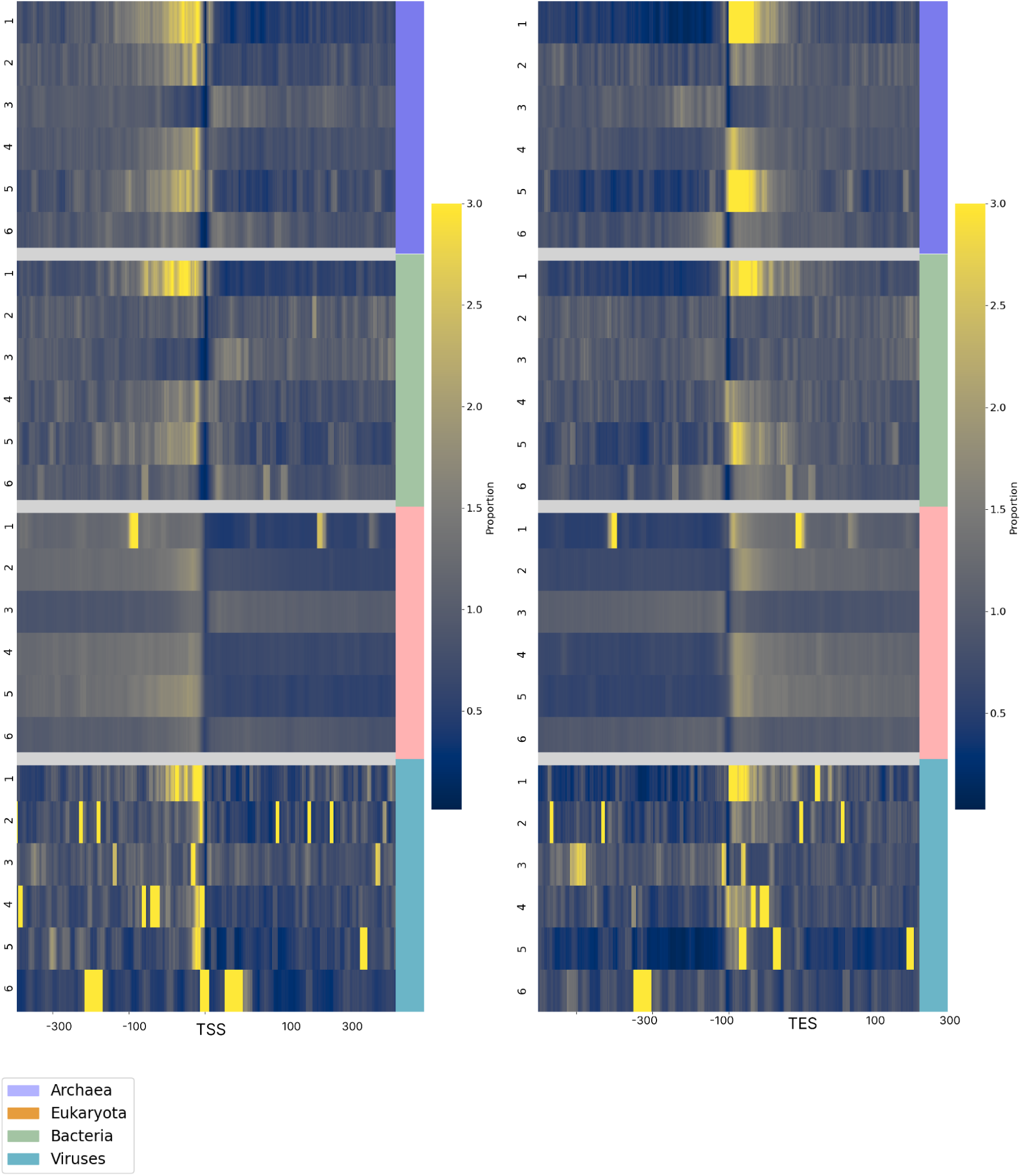
Distribution of STRs relative to TSSs/TESs in the three domains of life and viruses.

**Supplementary Figure 9:**
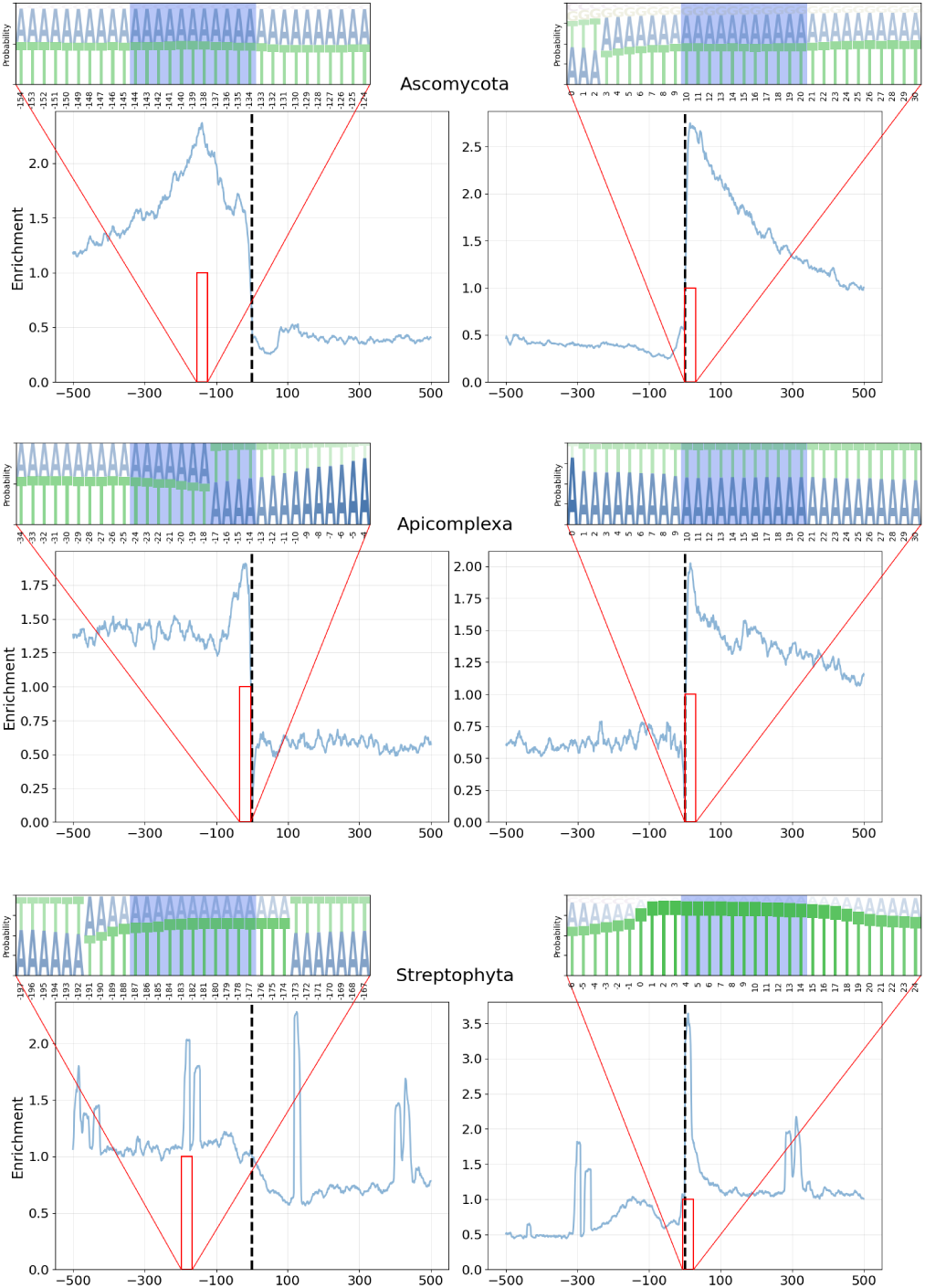
Distribution of mononucleotide STRs in Ascomycota, Apicomplexa and Streptophyta.

**Supplementary Figure 10:**
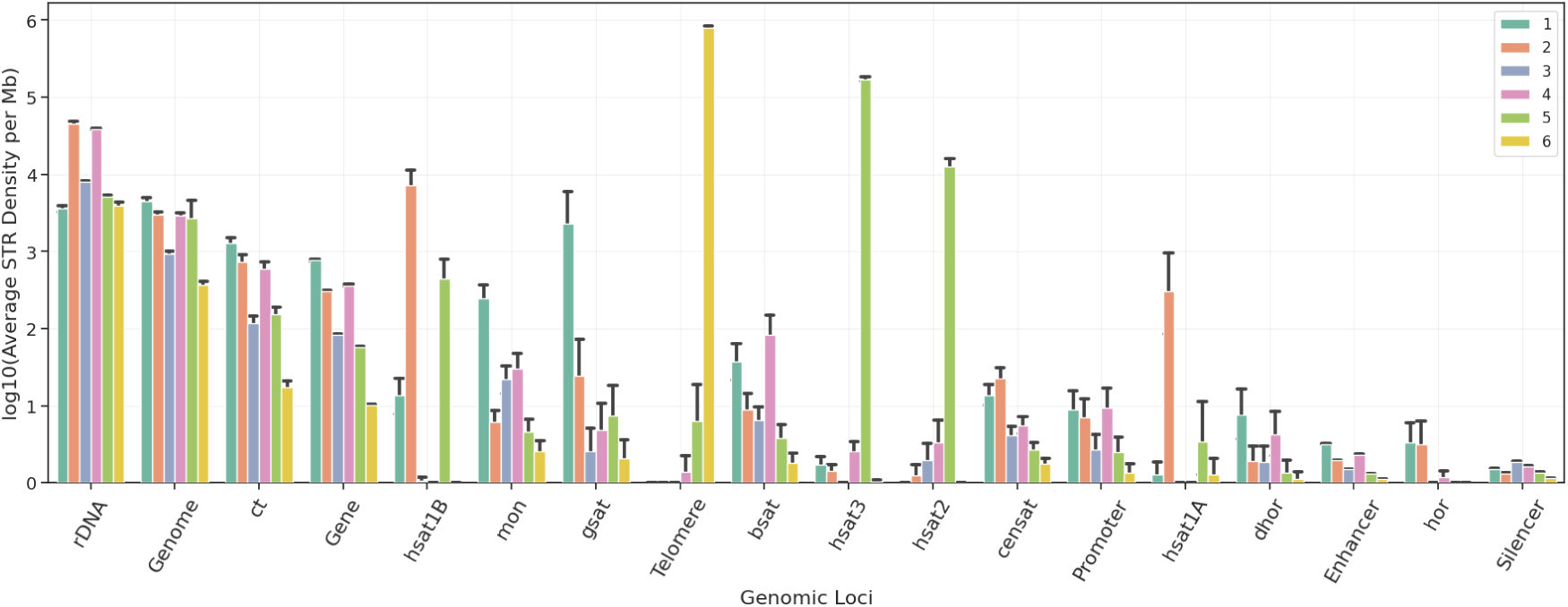
STR Density across human genome sub-compartments including centromeric repeats, separated by STR repeat length. Repeats include inactive αSat HOR (hor), divergent αSat HOR (dhor), monomeric αSat (mon), classical human satellite 1A (hsat1A), classical human satellite 1B (hsat1B), classical human satellite 2 (hsat2), classical human satellite 3 (hsat3), beta satellite (bsat), gamma satellite (gsat), other centromeric satellites (censat) and centromeric transition regions (ct).

**Supplementary Figure 11:**
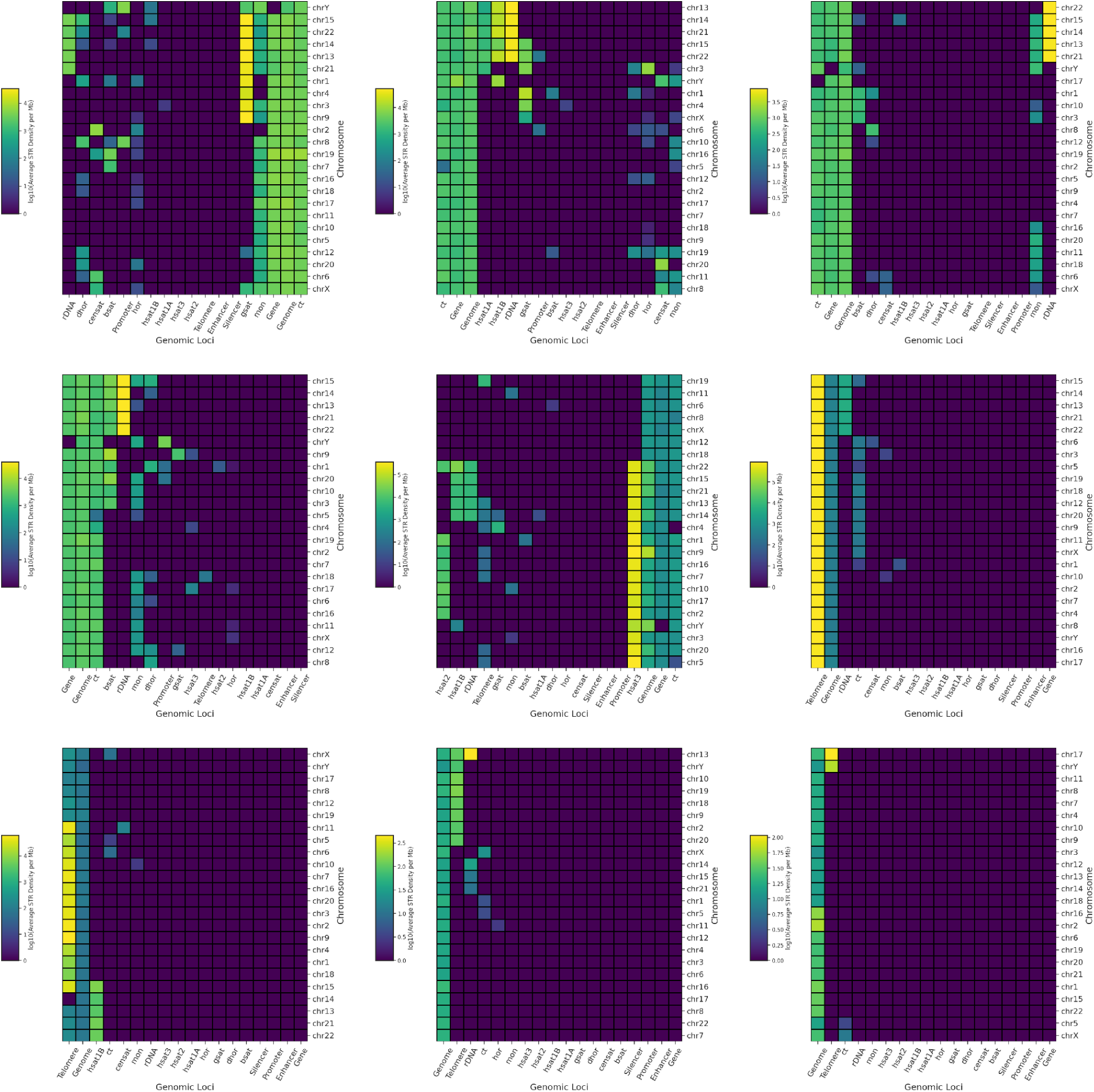
STR Density across the human genome sub-compartments separated by STR repeat length. Repeats include inactive αSat HOR (hor), divergent αSat HOR (dhor), monomeric αSat (mon), classical human satellite 1A (hsat1A), classical human satellite 1B (hsat1B), classical human satellite 2 (hsat2), classical human satellite 3 (hsat3), beta satellite (bsat), gamma satellite (gsat), other centromeric satellites (censat) and centromeric transition regions (ct). Results shown sequentially for one to nine bp STR units.

**Supplementary Figure 12:**
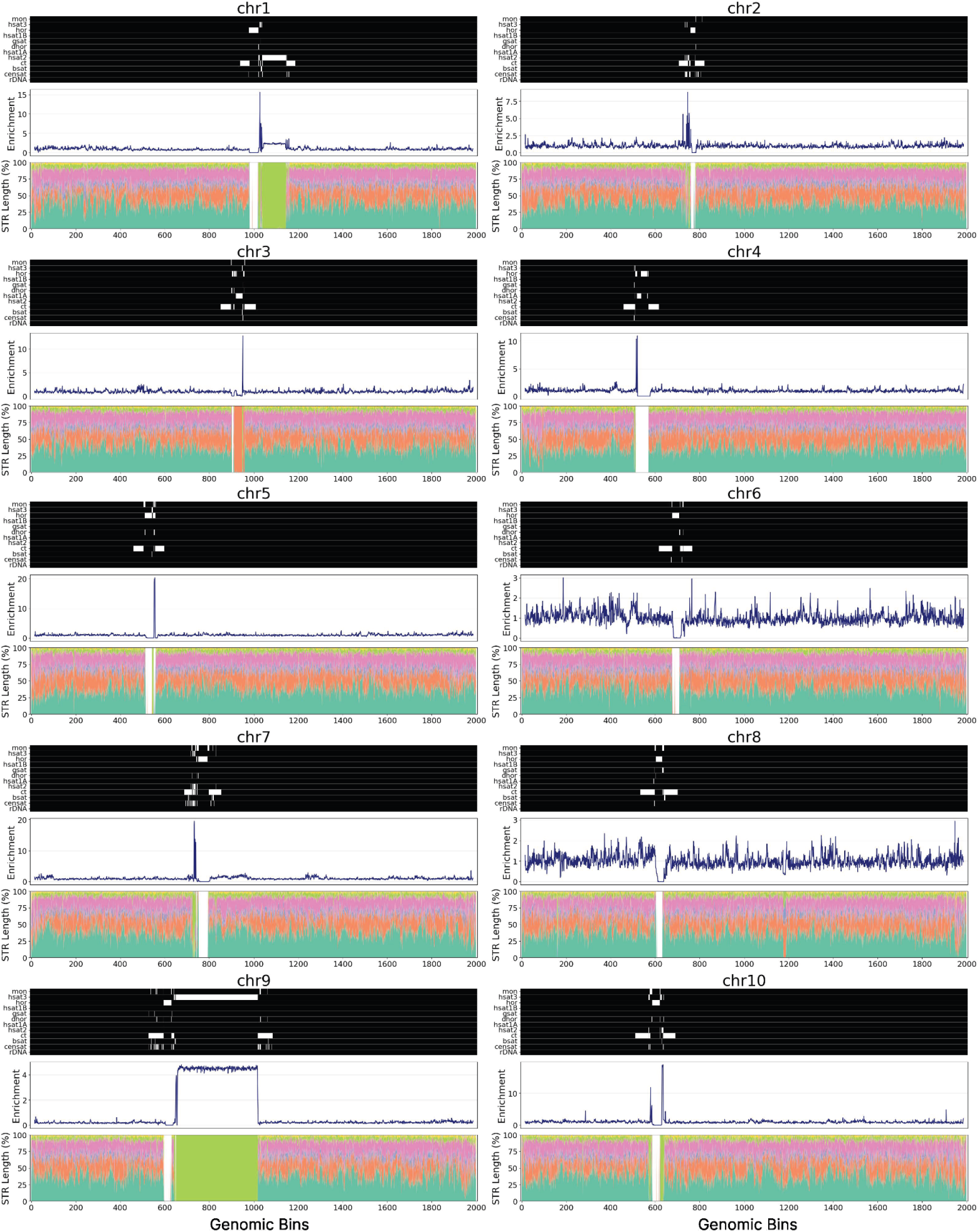
Characterization of STRs across chromosomes in the T2T reference human genome. Schematics show the distribution of STRs across different human chromosomes. The heatmap shows the different types of pericentromeric and centromeric repeats, with white color representing presence of the repeat in that genomic region. Line plots show the STR enrichment at each genomic bin for a chromosome. Stacked barplots show the results for different STR unit lengths.. Repeats marked in the heatmap include inactive αSat HOR (hor), divergent αSat HOR (dhor), monomeric αSat (mon), classical human satellite 1A (hsat1A), classical human satellite 1B (hsat1B), classical human satellite 2 (hsat2), classical human satellite 3 (hsat3), beta satellite (bsat), gamma satellite (gsat), other centromeric satellites (censat) and centromeric transition regions (ct).

**Supplementary Figure 13:**
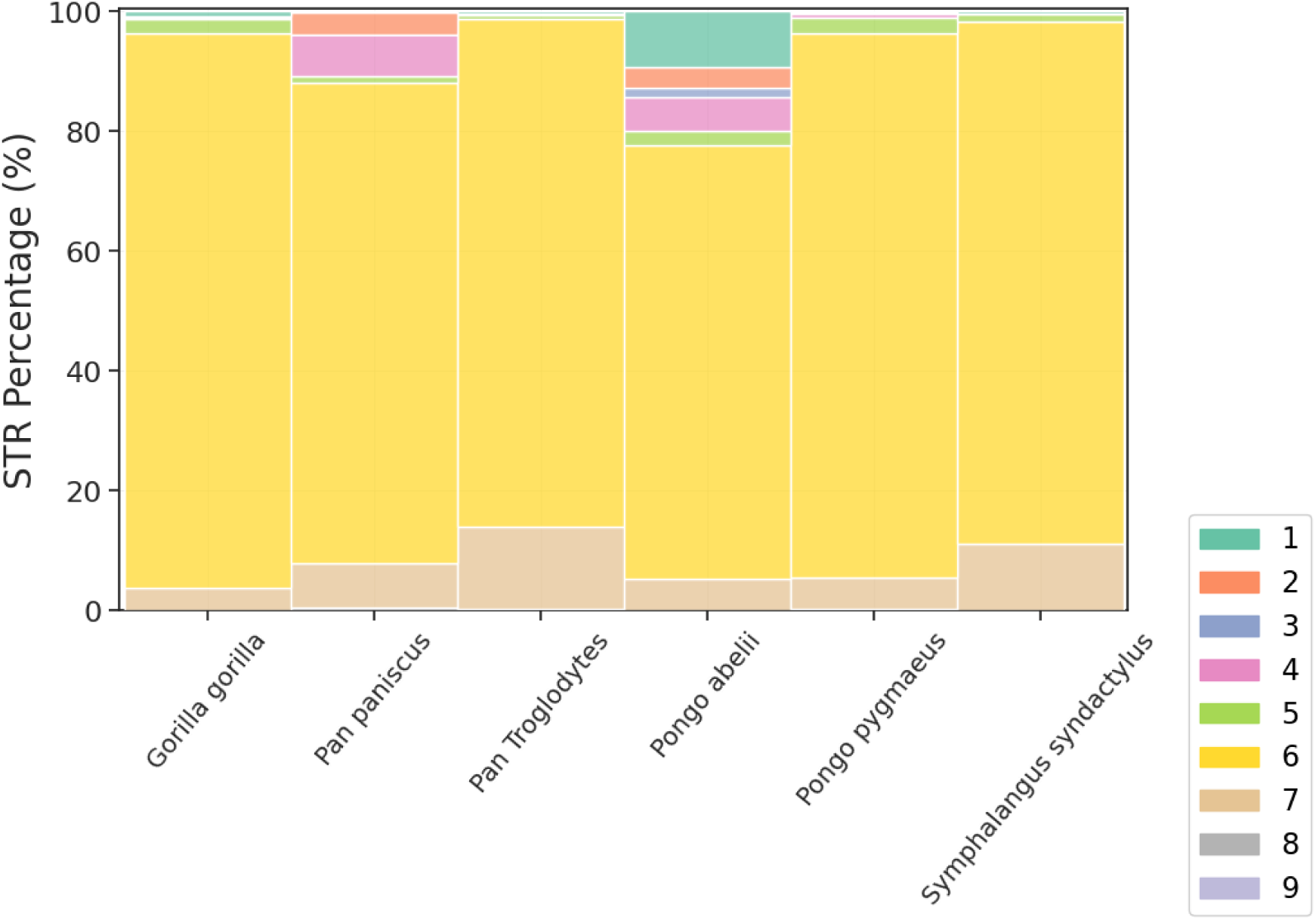
STR composition of telomeres for six primate species. Results shown for the STR unit lengths.

**Supplementary Figure 14:**
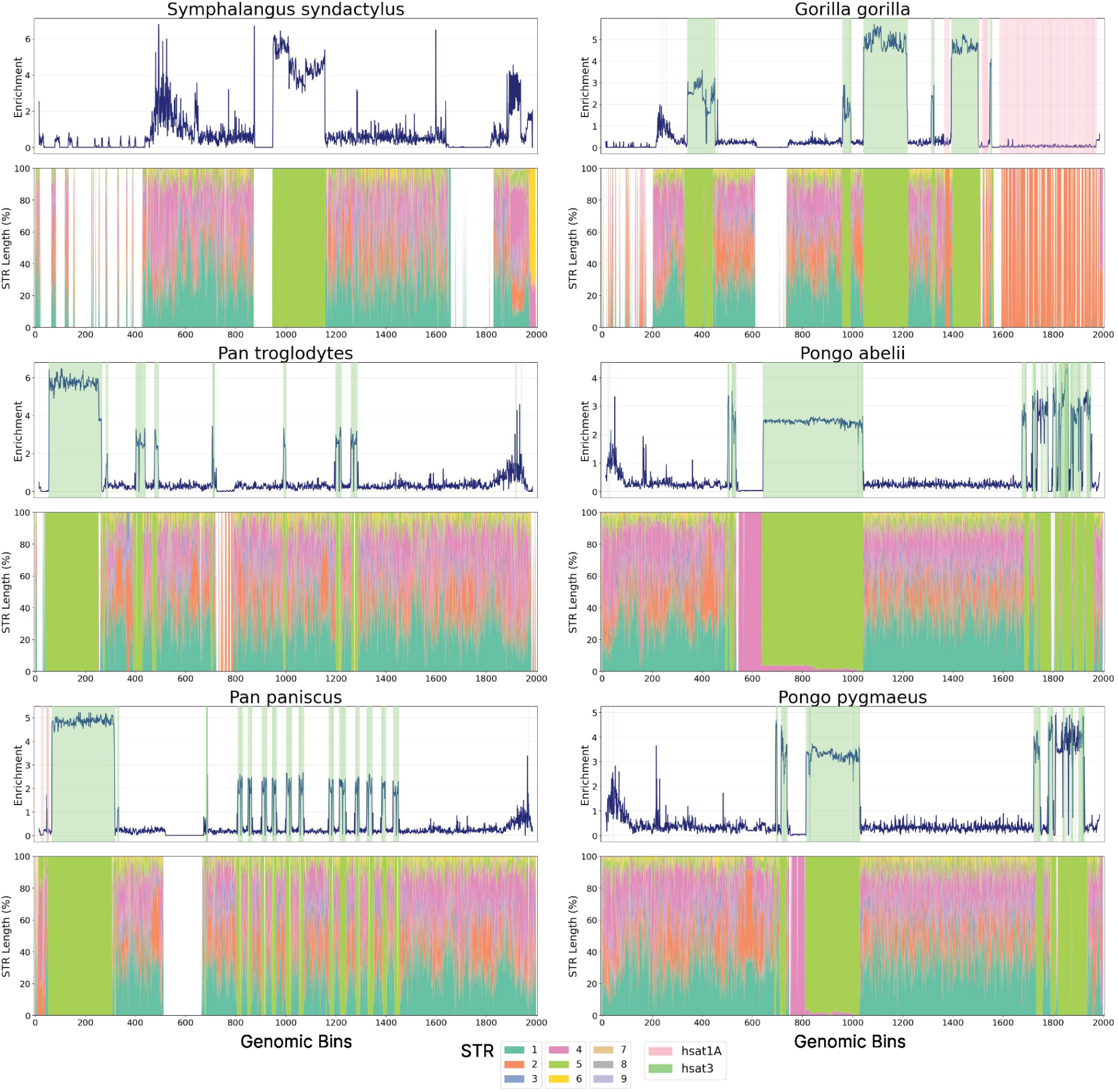
Characterization of STRs across the Y chromosomes of the T2T primate genomes. Highlighted in lightgreen and pink are the satellite array regions hsat3 and hsat1A, respectively, that include centromeric and centromeric repeats. Stacked barplots show the results for different STR unit lengths.

**Supplementary Figure 15:**
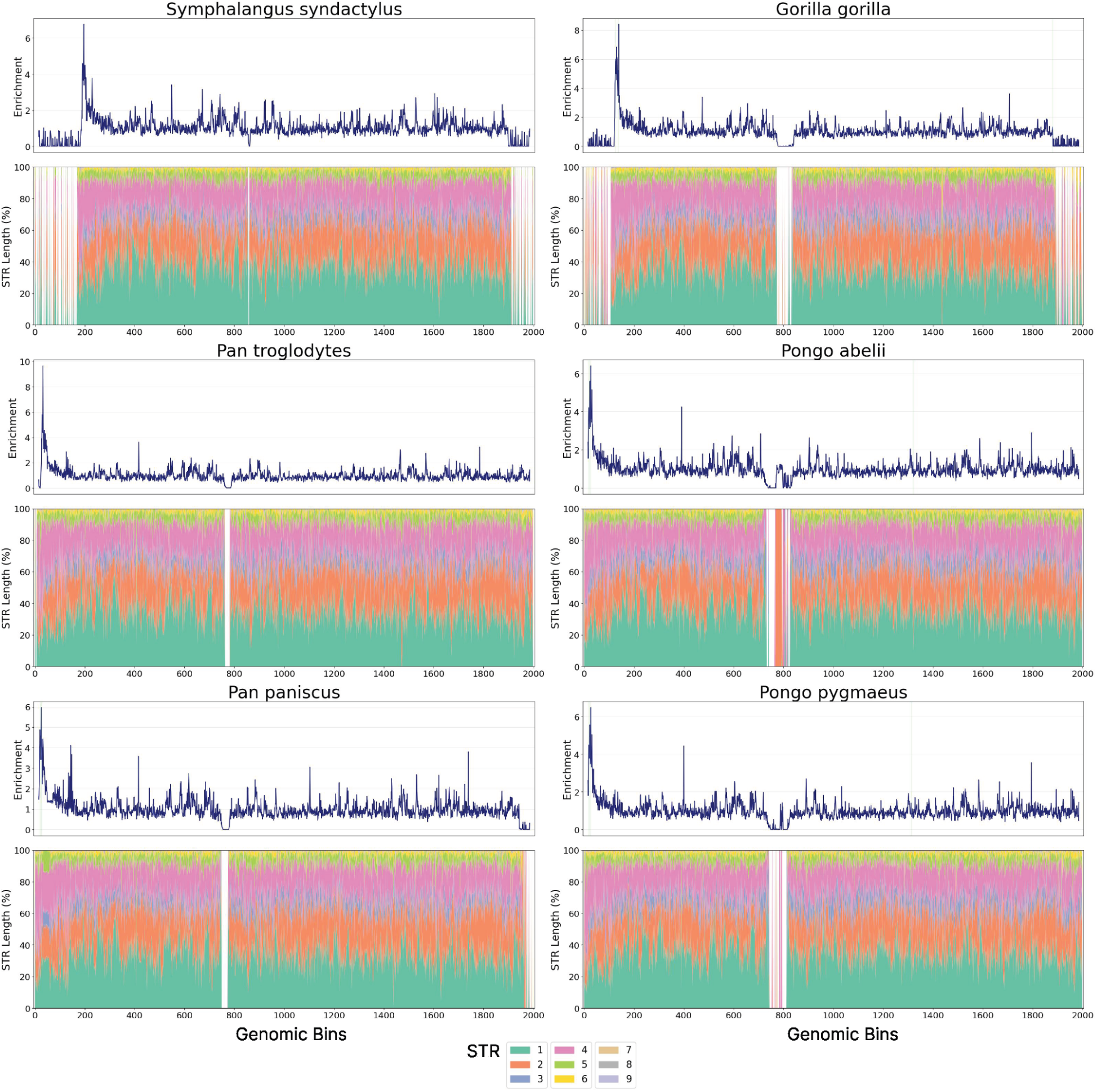
Characterization of STRs across the X chromosomes of the T2T primate genomes. Highlighted in pink are the satellite array regions that include centromeric and centromeric repeats. Stacked barplots show the results for different STR unit lengths.

**Supplementary Table 1:** Average STR Density per Mb for various GC-Thresholds.

| **GC Threshold** | **Fungi** | **Protista** | **Animalia** | **Plantae** | **Archaeabacteria** | **Eubacteria** | **Viruses** |
| --- | --- | --- | --- | --- | --- | --- | --- |
| 0% | 443,28 | 1425,12 | 964,04 | 886,14 | 75,07 | 81,21 | 132,49 |
| 10% | 260,75 | 492,27 | 535,73 | 477,48 | 60,46 | 62,22 | 94,04 |
| 20% | 257,00 | 483,46 | 529,99 | 471,60 | 59,96 | 61,74 | 92,64 |
| 30% | 227,41 | 442,48 | 419,82 | 427,53 | 55,23 | 57,33 | 83,47 |
| 40% | 177,79 | 375,23 | 369,22 | 371,28 | 49,48 | 51,67 | 66,88 |
| 50% | 168,92 | 370,72 | 273,59 | 360,53 | 49,03 | 51,18 | 66,06 |
| 60% | 99,39 | 166,45 | 63,76 | 224,54 | 40,85 | 45,91 | 46,94 |
| 70% | 48,23 | 92,62 | 37,79 | 104,37 | 21,76 | 34,43 | 28,53 |
| 80% | 36,08 | 64,32 | 23,82 | 85,58 | 15,72 | 30,54 | 23,27 |
| 90% | 30,02 | 56,99 | 20,68 | 74,08 | 14,04 | 29,00 | 21,88 |

**Supplementary Table 2:**
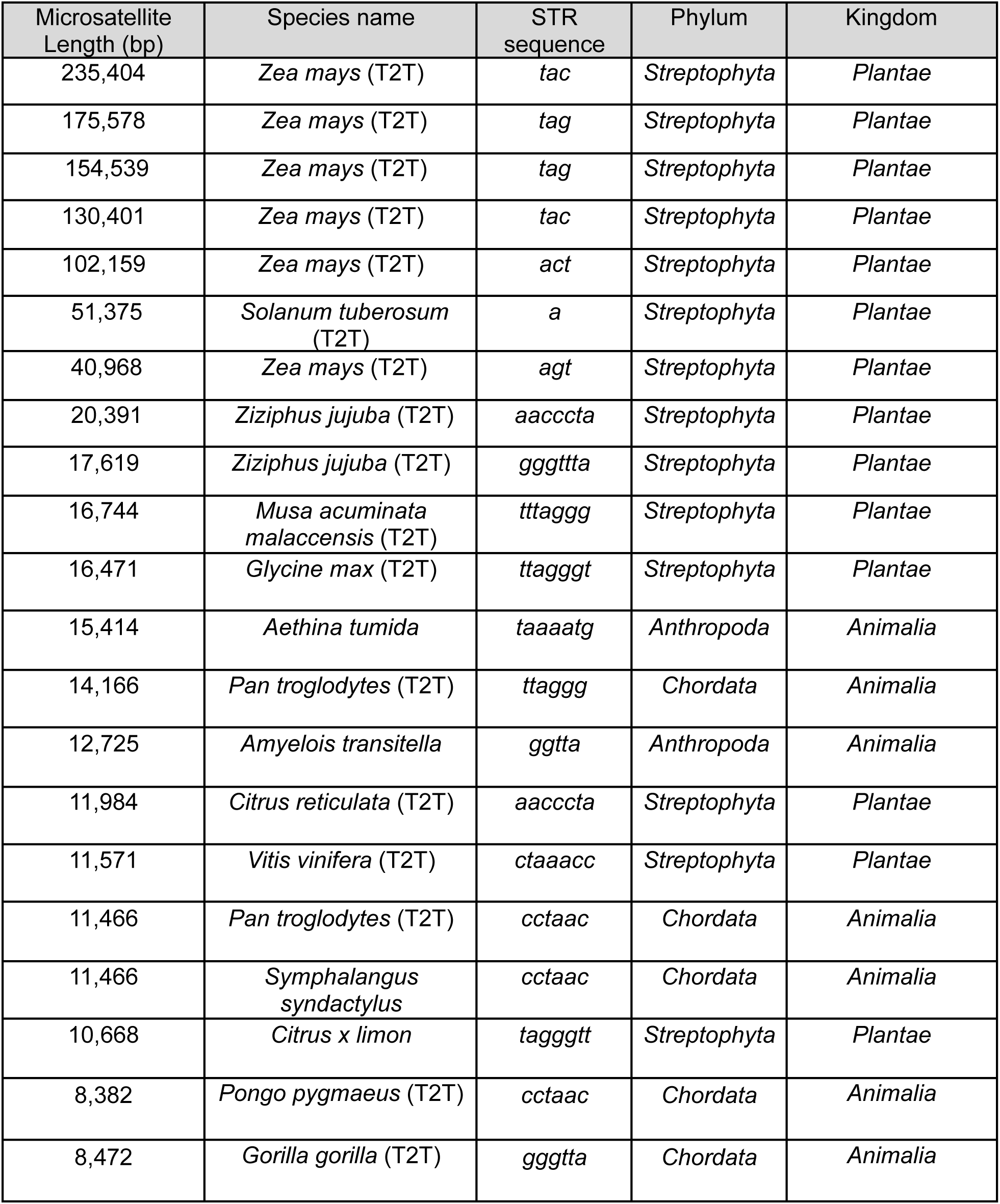
Longest, perfect STRs across the genomes studied. T2T genomes used are marked next to the species names.

